# Elevated synaptic PKA activity and abnormal striatal dopamine signaling in *Akap11* mutant mice, a genetic model of schizophrenia and bipolar disorder

**DOI:** 10.1101/2024.09.24.614783

**Authors:** Bryan J. Song, Yang Ge, Ally Nicolella, Min Jee Kwon, Bart Lodder, Kevin Bonanno, Antia Valle-Tojeiro, Nolan D. Hartley, Kira Perzel Mandell, John Adeleye, Deeksha Misri, Chuhan Geng, Sahana Natarajan, Inès Picard, Nate Shepard, Alyssa Hall, Jiawen Tian, Sameer Aryal, Zohreh Farsi, Xiao-Man Liu, Nader Morshed, Naeem M Nadaf, Horia Pribiag, Sean K Simmons, D. R. Mani, Beth Stevens, Prabhat S Kunwar, Zhanyan Fu, Evan Z. Macosko, Joshua Z. Levin, Bernardo L. Sabatini, Steven A. Carr, Borislav Dejanovic, Hasmik Keshishian, Adam J. Granger, Morgan Sheng

## Abstract

Loss-of-function mutations in *AKAP11* (a protein kinase A (PKA)-binding protein) greatly increase the risk of bipolar disorder and schizophrenia. To determine the neurobiological functions of AKAP11, we conduct multi-omic and neurobiological analyses of *Akap11* mutant mouse brains. We find that AKAP11 is a key regulator of PKA proteostasis in the brain whose loss leads to dramatically increased levels of PKA subunits and phosphorylated PKA substrates, especially in synapses. *Akap11* mutant mice show extensive transcriptomic changes throughout the brain, including prominent decreases in synapse-related genes sets. Gene expression is highly impacted in spiny projection neurons of the striatum, a brain region implicated in motivation, cognition and psychotic disorders. Real-time measurements of PKA activity reveal elevated basal PKA activity in the striatum of *Akap11^-/-^* mice, with exaggerated additional response to dopamine receptor antagonists. Behaviorally, *Akap11* mutant mice show abnormally prolonged locomotor response to amphetamine, deficits in associative learning and contextual discrimination, as well as depression-like behaviors. Our study connects molecular changes to circuit dysfunction and behavioral disturbance in a genetically valid animal model of psychotic disorder.

## Introduction

Bipolar disorder (BD) and schizophrenia (SCZ) are debilitating psychiatric conditions within the psychosis spectrum whose biological basis remains unknown. Bipolar disorder is typically characterized by episodes of mania alternating with periods of depression. Schizophrenia is characterized by positive symptoms including hallucinations and delusions that occur alongside negative symptoms that include emotional impairment, and is often associated with cognitive symptoms. Together, BD and SCZ account for a substantial fraction of the global mental health crisis: the lifetime prevalence of BD is 1-2%^1^ and SCZ is approximately 0.7%^2^. Both diseases have substantial heritability (60-80%)^3,4^, suggesting that gene-derived molecular programs contribute towards disease risk.

Recent landmark exome sequencing efforts have led to the discovery of several genes whose loss of function greatly increase the risk for schizophrenia (SCZ)^5^ and bipolar disorder (BD)^6^. These rare variant mutations are associated with large odds ratios (3-50), conferring much more substantial risk than common variants previously identified by genome-wide association studies (odds ratios <1.1)^7,8^. In further contrast to GWAS-associated loci, these mutations occur in coding sequences of specific genes and can be inferred to be loss of function in nature.

Mutations in *AKAP11* (A-kinase anchoring protein 11) are associated with greatly elevated risk of both SCZ and BD^5,6,9,10^. *AKAP11* has not been associated with autism spectrum disorder, epilepsy or intellectual disability^6^ and therefore is a relatively specific risk gene for the SCZ-BD psychotic disorder spectrum. In contrast, previously studied large-effect SCZ risk variants^11–13^ are also associated with intellectual disability and developmental disorders^5^, making *Akap11* animal models particularly interesting to study. In our previous neurophysiologic study, we reported that *Akap11* loss-of-function mutant mice recapitulate several electroencephalogram (EEG) abnormalities found in schizophrenic patients, including increased power of resting gamma oscillations, non-rapid eye movement (NREM) sleep reduction, and gene-dose dependent reduction in sleep spindles^14^. The molecular, cellular and circuit mechanisms by which *Akap11* deficiency results in brain dysfunction have not been studied.

The *Akap11* gene encodes a member of the A-kinase anchoring proteins (AKAPs), which bind Protein Kinase A (PKA) and are typically localized to compartments within a cell, targeting PKA to specific subcellular domains^15^. In non-neuronal cell lines, AKAP11 is reported to localize to punctate structures, including multivesicular bodies^16^ and peroxisomes^17^. In addition to PKA, AKAP11 is reported to interact biochemically with the protein kinase GSK3β^18^, Protein Phosphatase 1^19^, IQGAP1^20^, GABA_C_ receptors^21^, and vesicle-associated proteins A and B (VAPA, VAPB)^22^, each of which is implicated in signaling and trafficking pathways that influence synaptic plasticity and neuronal development. Intriguingly, AKAP11 not only binds but also mediates inhibition of GSK3β^18^, a target of the mood-stabilizing drug lithium that is used to treat BD. AKAP11 also functions as an autophagy receptor for the regulatory subunit 1 of PKA (PKA R1α) which promotes protein-level degradation of PKA R1α in the context of cellular stress^23^.

Because AKAP11 has not been studied in the brain, there is insufficient knowledge to bridge the cell biology of *AKAP11* and psychiatric disease risk. In this study, we use unbiased multi-omic approaches to explore the consequences of *Akap11* loss-of-function across widespread brain regions and cell types, spanning several postnatal ages and within synaptic compartments. To follow up on multi-omic analyses, we carry out targeted real-time measurements of net striatal PKA activity in the context of dopaminergic signaling using *in vivo* fiber photometry. Our data uncover a fundamental role of *AKAP11* in synapse function and highlight the striatum as a nexus of circuit dysfunction in a model of psychosis risk with human-genetics validity.

## Results

### *Akap11* mutant mice exhibit activity, cognitive and affective behavioral phenotypes

To investigate the effects of *Akap11* deficiency on mouse behavior, we conducted a multi-level set of behavioral experiments encompassing long-term naturalistic observations in the home cage, machine learning based analysis of complex motor behavior, and specific tests that may be relevant to psychiatric symptom domains. We first examined the general activity patterns of *Akap11^+/-^, Akap11^-/-^* and wild-type (WT) littermate controls continuously in their home cages from 4 weeks to 16 weeks of age using an automated home cage activity monitoring system^24–26^. *Akap11* mutants were less active than WT littermates during their night-time (active) and daytime (resting) periods, in a gene-dose dependent manner (Fig. 1a,b). This hypoactivity was apparent at 4 weeks of age (at the beginning of monitoring) and remained until 12 weeks (Fig. 1c). When a running wheel was introduced for 1 week during the assay, *Akap11^-/-^* mutant mice showed less engagement than WT^27^ (Fig. 1d).

**Figure 1:**
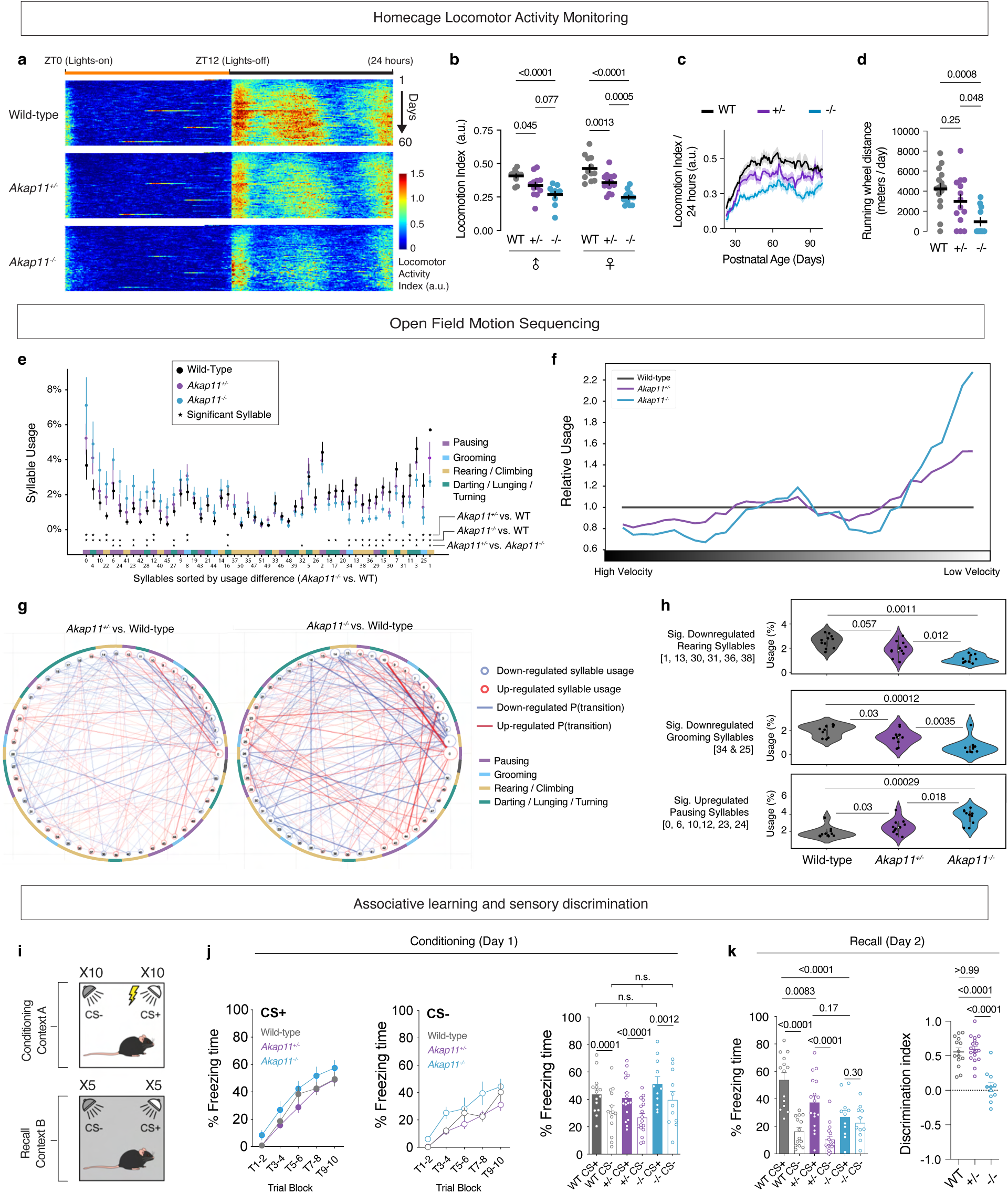
Decreased locomotion, associative learning and sensory discrimination in *Akap11* mutant mice. **a**, Heatmap of average activity of each genotype across 24 hours in a homecage locomotor activity monitoring system. Each row is 24 hours, and 60 rows representing 60 consecutive days are shown. Yellow and black bars indicate lights-on and lights-off periods respectively. ZT (Zeitgeber Time) refers to the number of hours past the lights-on time in light-dark scheduled conditions. **b**, Averaged locomotor activity of each WT and *Akap11* mutant mice, across 60 days of the assay. **c**, Locomotor activity averaged across 24 hours, aligned according to animals’ postnatal ages. **d**, Running wheel usage during a 1 week test period. **e**, Syllable usage of WT and *Akap11* mutant mice (n=11 mice for WT, n=11 mice for *Akap11^+/-^, and* n=10 mice for *Akap11^-/-^*), sorted by usage difference between WT and *Akap11^-/-^*. Color codes on the x-axis represent annotated behaviors. * Represents represent syllables with FDR <0.1. Data are represented as mean +/- 95% confidence interval. **f**, Relative syllable usage of WT and Akap11 mutant mice sorted by velocity. **g**, Syllable transition matrix showing the upregulated (red lines) and downregulated (blue lines) transition probability between *Akap11^+/-^* and WT (left) or between *Akap11^-/-^* and WT (right). **h**, Average usage of significantly downregulated rearing syllables (top), grooming syllables (middle), and significantly upregulated pausing syllables (bottom) between *Akap11* mutants and WT (n=11 mice for WT, n=11 mice for *Akap11^+/-^, and* n=10 mice for *Akap11^-/-^*). The box plots display the median (white dot) and interquartile range (25^th^-75^th^ percentile); the whiskers represent upper and lower adjacent values; the outer bounds represent the data distribution. **i**, Schematic diagram of the associative learning and sensory discrimination task. **j**, During fear conditioning (Day 1), freezing time (%) to CS+ (left) and CS- (middle) by WT and *Akap11* mutant mice. Bar graph (right) represents the freezing time (%) during trials 6-10. **k**, During fear recall (day 2), freezing time (%) and to CS+ and CS- by WT and Akap11 mutant mice (left). Discrimination index (right) is defined as: (CS+ freeze - CS- freeze) / (CS+ freeze + CS- freeze). **b,c,d,j,k**, data are represented as mean +/- SEM, and bar graphs as mean + SEM. **b,d,h,** One-way ANOVA with Tukey’s post hoc test. See Supplementary Data 1 for statistics. **j,k,** Repeated measures two-way ANOVA with Tukey’s post hoc test for CS+ and CS- freezing time (%), and one-way ANOVA with Bonferroni post hoc test for discrimination index. See Supplementary Data 1 for statistics.

To examine volitional (“exploratory”) behavior at finer resolution, we applied motion sequencing (MoSeq), an unsupervised machine learning based tool that decomposes continuous 3D video recordings into discrete, repeated behavioral motifs (“syllables”)^28,29^. During a 30-minute open field trial, *Akap11^-/-^* mice displayed significant differences from WT littermates (*P* < 0.05) in their usage of a wide range of behavioral syllables (26 of 51 syllables) (Fig. 1e, Supplementary Fig. 1a) including those related to locomotion, pausing, grooming and rearing (Fig. 1e,h). Many of these same syllables (10 of 26 syllables significant in *Akap11^-/-^*) were differentially utilized in *Akap11^+/-^* mice but to an intermediate degree. Gene-dose dependent alterations in syllable usage were also apparent in the proportions of low/high velocity syllables (Fig. 1f) and transition probabilities between syllables (Fig. 1g). *Akap11* mice were hypolocomotive in home cage and MoSeq assays, consistent with prior open-field measurements^14,30^. In the rotarod test, *Akap11* mutant mice did not differ from WT in latency to fall, suggesting that their low activity was not due to grossly impaired motor coordination (Supplementary Fig. 1b).

We next measured mouse behaviors that are commonly used as models of positive and negative symptoms^31^. *Akap11* mutant mice performed similarly to WT in prepulse inhibition (Supplementary Fig. 1c), suggesting intact sensory gating. *Akap11* mutant mice also had normal social behaviors – their social preference to a novel mouse over an object (Supplementary Fig. 1d), or preference to a novel mouse versus a familiar mouse (Supplementary Fig. 1e), did not differ from WT controls. In the forced swim test (often used to infer “depressive-like” states^32^), *Akap11* mutant mice displayed greater immobility in a gene-dose dependent fashion (Supplementary Fig. 1f). *Akap11* mutants performed normally in the sucrose preference test if they were tested as a “naïve” cohort; interestingly, however, if the sucrose preference test followed a forced swim test on the preceding day, then *Akap11^-/-^* mutants showed robustly impaired sucrose preference compared with WT controls (Supplementary Fig. 1g), a pattern that could be interpreted as anhedonia^33^. Together with the greater immobility in the forced swim test, this pattern could be interpreted as depressive-like and anhedonia-like behavior that is induced by prior stress^33^.

Cognitive deficits are highly prevalent in schizophrenia, including attention deficits, memory impairments, and reduced executive control^34^. We designed a sensory discrimination task using auditory fear conditioning to simultaneously test *Akap11* mutants for associative learning and sensory / context discrimination (Fig. 1i). Mice were presented with a certain tone (Conditioned Stimulus + [CS+]) that is associated with footshock (Unconditioned Stimulus, US) and a different tone (Conditioned Stimulus -, CS-) that is not associated with any US. During the conditioning day (Day 1), WT, *Akap11^+/-^*, and *Akap11^-/-^*mice all displayed increased freezing during CS+ compared with during CS-, indicating successful learning of fear conditioning and CS discrimination (Fig. 1j). During the recall test (Day 2), *Akap11^+/-^* and *Akap11^-/-^* had impaired fear recall as measured by less freezing time in response to CS+ compared with WT (Fig. 1k, left). While WT and *Akap11^+/-^* mice showed increased freezing to CS+ than to CS-, *Akap11^-/-^*mice froze similarly to both CS+ and CS- (Fig. 1k, right). Thus *Akap11^+/-^* and *Akap11*^-/-^ mutant mice had gene-dose dependent impairment in fear conditioning, whereas *Akap11^-/-^* mice also exhibited impaired ability to discriminate CS+ and CS- one day after conditioning, suggesting a generalization of fear memory in the homozygous mutant. Together, these data show behavioral abnormalities in *Akap11* mutant mice in domains (mood, cognition) that are relevant to neuropsychiatric disease.

### Identifying the AKAP11 interactome in brain

Little is known about AKAP11 in the brain, so we characterized the protein interactome of AKAP11 by immunoprecipitation of AKAP11 from cortical extracts followed by mass spectrometry (coIP-MS, Fig. 2a,b; Supplementary Fig. 2a). Of the ∼2000 proteins identified by mass spectrometry, 68 proteins were enriched (*P* < 0.05) in AKAP11 coIP from *Akap11* WT cerebral cortex versus coIP from negative control *Akap11^-/-^* (knockout) mice^30^; these were considered specific interactors of AKAP11 (Fig. 2c; Supplementary Data 2). This approach yielded putative AKAP11-associated proteins that were largely overlapping with those found by comparison of AKAP11 coIP vs control IgG coIP from WT brains (Supplementary Fig. 2b). Of the 68 proteins that were specifically associated with AKAP11 in the brain, only 7 were previously reported in cell lines or other tissues, including PKA subunits, GSK3β, and VAPA and VAPB (Fig. 2d)^18,22^. The biochemical association of AKAP11 with PKA catalytic and regulatory subunits and several other interactors (VAPA, VAPB, GSK3β, DRYK1A) was confirmed by coIP-immunoblot experiments (Fig. 2e-j). AKAP11 did not associate with the cytoskeleton protein Vinculin, indicating specificity for the above results (Fig. 2e).

**Figure 2:**
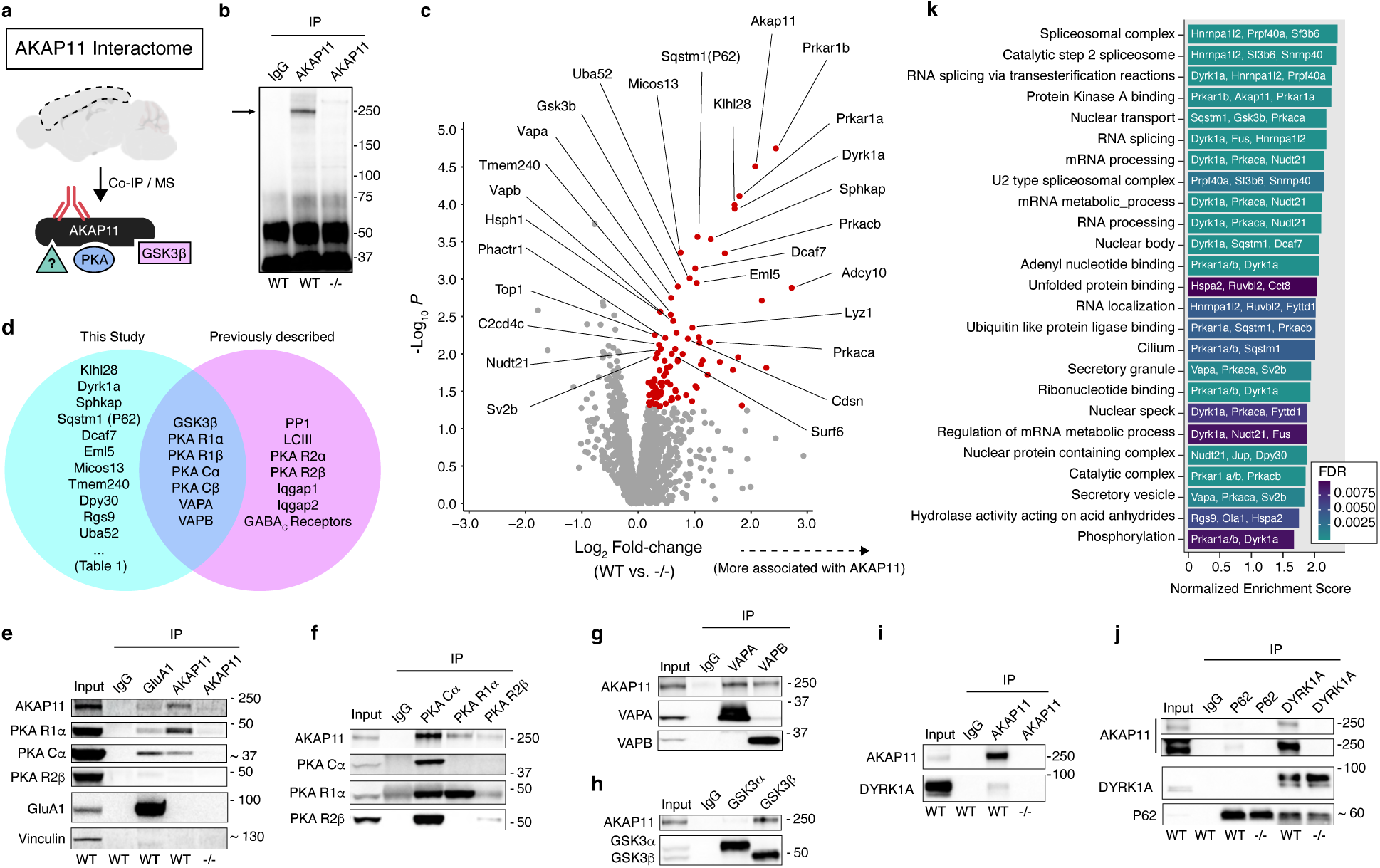
Characterization of the AKAP11 interactome in the brain. **a,** Schematic of AKAP11 immunoprecipitation (IP) and its most notable, known interactors. **b,** Immunoblot of AKAP11 in IPs of anti-AKAP11 and control IgG in 12-week-old (12wk) wild-type (WT) or *Akap11*^-/-^ cortical tissue. This experiment was repeated in >5 independent experiments, with similar results. **c,** Volcano plot comparing Log_2_ Fold-Change (Log_2_FC) against *P* value for AKAP11 IP in WT vs *Akap11*^-/-^, displaying all proteins detected by coIP-MS. A moderated two-sample t-test was applied to the sample group datasets. Proteins that were enriched (red) by anti-AKAP11 IP from WT versus *Akap11^-/-^* cortical extracts. Enriched proteins, defined by having a nominal *P* value < 0.05 and Log_2_FC > 0, are colored red. Downregulated proteins are mostly antibody-related artifacts and can be found in Supplementary Data 2. **d,** Venn diagram comparing previously published AKAP11 interactions with the brain interactome in this study. Full results are displayed in Supplementary Data 1. **e,** Immunoblot of AKAP11, PKA subunits and control proteins, in IPs of AKAP11, GluA1 or IgG control in WT or *Akap11^-/-^* cortical lysate. This experiment was repeated in >3 independent experiments, with similar results. **f,** Immunoblot of AKAP11 in IPs of PKA subunits, in WT cortical lysates. This experiment was repeated in >2 independent experiments, with similar results. **g-j,** Immunoblot with indicated antibodies, in IPs of anti-GSK3α, GSK3β, VAPA, VAPB, AKAP11, P62 or DYRK1a in WT or *Akap11*^-/-^ cortical lysate. These experiments were repeated in at least 2 independent experiments, with similar results. **k,** Gene set enrichment analysis of all proteins displayed in 1c, showing selected gene ontologies with a positive normalized enrichment score (NES) and adjusted *P*-value (FDR) < 0.01. Redundant (similar) pathways were removed for clarity. The top most enriched proteins within each ontology are listed within the histograms. GSEA uses a Kolmogorov-Smirnov test, two tailed, with Benjamini-Hochberg (B-H) False Discovery Rate (FDR) correction applied.

As expected for an AKAP protein^15^, we observed high enrichment of multiple subunits of PKA in AKAP11 immunoprecipitates (Fig. 2c,e). PKA exists as a tetrameric holoenzyme with regulatory subunits that bind and suppress the kinase activity of catalytic subunits. In AKAP11 coIPs, there was greater degree of enrichment for regulatory subunits PKA R1α and PKA R1β (encoded by *Prkar1a* and *Prkar1b*) than the catalytic subunits PKA Cα, PKA Cβ (encoded by *Prkaca* and *Prkacb*), which is consistent with AKAP11 directly binding to regulatory subunits of PKA^35^. Adenylyl cyclase 10 (encoded by *Adcy10*) was also enriched, and this enzyme produces cyclic AMP, which binds and activates PKA. There was only modest enrichment for type 2 regulatory subunits (PKA R2α, PKA R2β. Supplementary Fig. 2c) even though AKAP11 has high affinity binding sites for R2^35^. We confirmed that AKAP11 also associates with GSK3β in the brain, consistent with prior studies in cell lines^18,35,36^ and did not detect an association with GSK3α (Fig. 2h).

Our coIP-MS experiments from brain uncovered multiple novel interactors of AKAP11 including DYRK1A, SPHKAP, Synaptic Vesicle Protein 2B (SV2B), and P62 (also known as SQSTM1) (Fig. 2c,d). By coIP-immunoblot, we confirmed AKAP11 binding to DYRK1A (Fig. 2i,j), a tyrosine kinase that is genetically linked to autism spectrum disorders^37^ and is implicated in neurodevelopment, neurogenesis and neurite outgrowth^38–41^. Association with VAMP-associated Proteins A and B (VAPA/VAPB) and SPHKAP is suggestive of localization near endoplasmic reticulum (ER). VAPA/B are ER-resident proteins that facilitate inter-organelle contact and vesicular transport and SPHKAP interacts with VAP proteins to regulate calcium signaling from ER stores^42^. We also explored the AKAP11 interactome by coIP from the brains of *Akap11^+/-^* (heterozygote) mutants, which mimics the human heterozygous genotype that is associated with greatly increased risk for SCZ/BD^6^. Proteins coIP’d with AKAP11 were largely similar in WT and *Akap11^+/-^* (Supplementary Fig. 2d,e).

To gain further insight into molecular pathways in which AKAP11 might play a role, we conducted gene-set enrichment analysis (GSEA)^43^ on AKAP11-associated proteins (Fig. 2k, Supplementary Data 3). In addition to expected gene ontology (GO) pathways like “Phosphorylation”, we observed enrichment in AKAP11 coIP of the “Ubiquitin like protein ligase binding” GO term, which includes the selective autophagy receptor P62 and RPL40 (encoded by *Uba52*) (Supplementary Fig. 2f). In cell lines, AKAP11 interacts with autophagosomes, intracellular vesicular structures involved in protein degradation^44^, through the selective autophagy receptor LC3^23,45^. In our coIP experiments, we found an association of AKAP11 with P62, but not LC3 (Supplementary Fig. 2c), suggesting that in the brain, AKAP11 also interacts with autophagosomes but possibly by distinct molecular mechanisms.

### Accumulation of PKA in *Akap11* mutant mice

Given our finding that AKAP11 could interact with autophagosomes, coupled with previous studies showing its role in targeting proteins for degradation, we tested how the abundance of AKAP11-interacting proteins was altered by loss of AKAP11. By immunoblotting of total cortical lysate, we found a striking elevation of PKA protein levels (regulatory (∼6x) and catalytic (∼9x) subunits in *Akap11^-/-^* mice) (Fig. 3a, Supplementary Fig. 3a). The association of AKAP11 with P62 in the brain (Fig. 2c) suggests the possibility that AKAP11 regulates degradation of PKA by receptor-mediated selective autophagy. Levels of other AKAP11 interactors (GSK3β, VAPA/B, P62 and DYRK1A) were not substantially altered in *Akap11* mutant brains.

**Figure 3.**
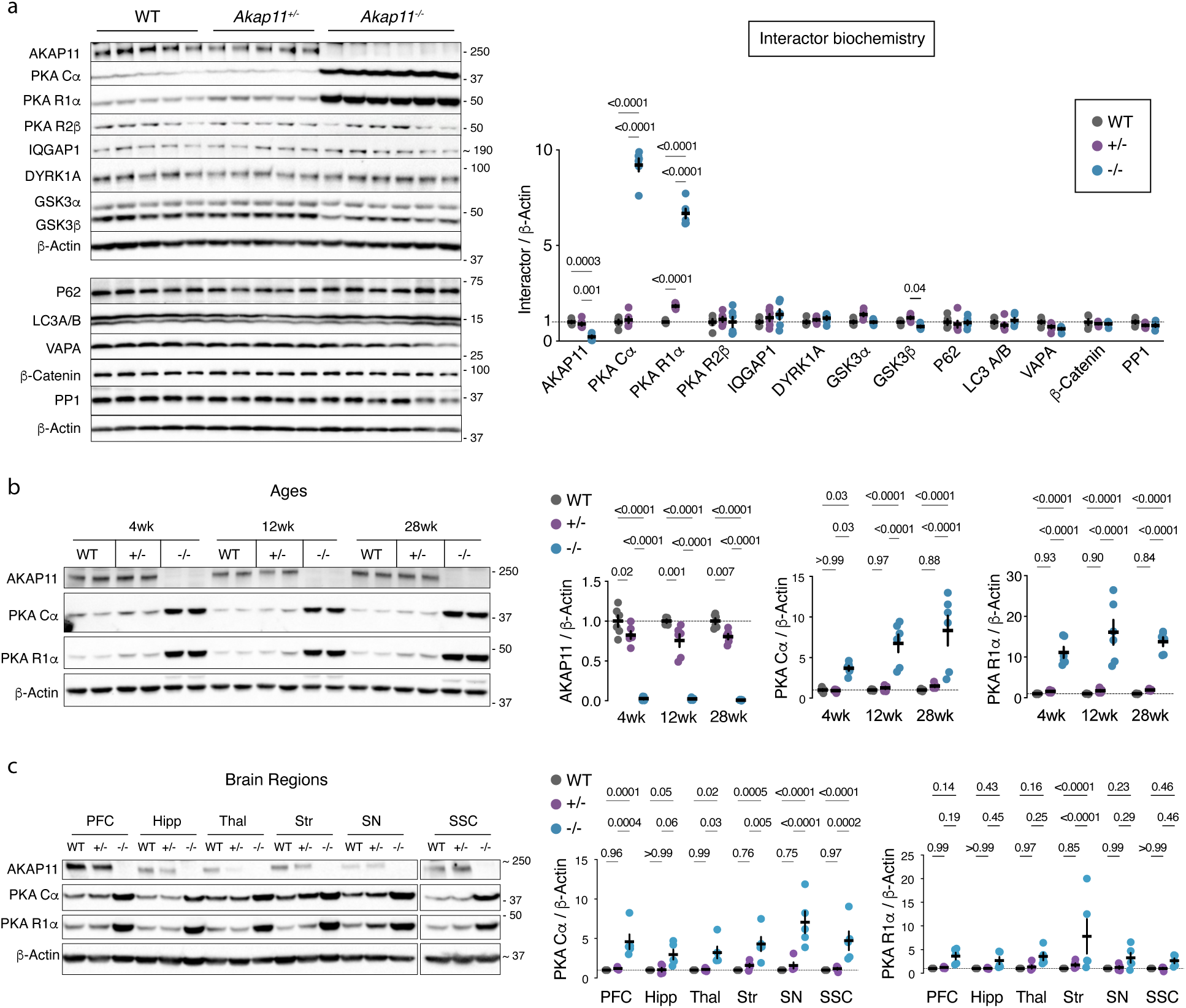
PKA protein levels are elevated across brain regions and ages of *Akap11* mutant mice. **a**, Immunoblots and quantifications of 12wk cortex total lysate using antibodies against AKAP11-interacting proteins. Measurements are normalized to β-Actin and to WT controls for each antibody. *P* < 0.05 comparisons are shown. **b,** Immunoblots and quantifications of 4wk, 12wk and 28wk cortex total lysate using antibodies against AKAP11 and PKA subunits. Measurements are normalized to β-Actin and to WT controls within an age group. **c,** Immunoblots and quantifications from microdissected tissue total lysate from 4wk mice using antibodies against AKAP11 and PKA subunits. Measurements are normalized to β-Actin and to WT controls within a region. **a-c,** Data are represented as mean +/- SEM. Two-way ANOVA with Tukey’s post hoc test. See Supplementary Data 1 for exact *P*-values, statistical tests and sample sizes for each comparison.

The marked elevation of PKA subunits in *Akap11^-/-^* mice was apparent at all ages examined (4, 12, 28 weeks of age, Fig. 3b). PKA subunits were increased not only in the cortex, but across all brain regions examined: prefrontal cortex (PFC), somatosensory cortex (SSC), striatum (Str), hippocampus (Hipp), thalamus (Thal) and substantia nigra (SN) (Fig. 3c). Compared with the multiple-fold increases in the homozygous (*Akap11^-/-^*) mutant, changes in PKA protein levels in the heterozygous (*Akap11^+/-^*) mutant brain were much more modest (Fig. 3a-c). Nonetheless, comparison with WT animals showed that PKA R1α and Cα subunits were consistently elevated by 20%-75% in *Akap11^+/-^* (Supplementary Fig. 3b,c). In the course of these immunoblotting experiments, we noted that levels of AKAP11 protein itself were abolished in the homozygous mutant (*Akap11^-/-^*) but minimally reduced in heterozygous (*Akap11^+/-^*) mutants (Fig. 3a,b, Supplementary Fig. 3a,d), despite gene-dose dependent reduction of *Akap11* RNA (Supplementary Fig. 3e). This pattern is suggestive of post-transcriptional homeostatic compensation of *Akap11* expression when just one copy of the *Akap11* gene is deleted, and might explain the relatively modest effect on PKA in *Akap11^+/-^*vs. *Akap11^-/-^*. Quantitative immunofluorescence imaging of PKA R1α protein in brain sections showed increased intensity across all brain regions of *Akap11^-/-^* mice (Supplementary Fig. 3f). Level of *PKA R1α* RNA was not significantly altered in *Akap11^-/-^* (Supplementary Fig. 3g), suggesting that AKAP11 regulates the stability or turnover of PKA at the protein level.

### Protein changes and elevated PKA at synapses of AKAP11 deficient mice

PKA plays a key role in regulation of synaptic plasticity^46^. To understand the consequences of *Akap11* deficiency and PKA elevation in synapses, we investigated protein changes in purified synaptic (“PSD (postsynaptic density)”) fractions of *Akap11* mutant mice by quantitative mass spectrometry-based proteomics and phosphoproteomics using Tandem Mass Tag (TMT) labeling (see methods). Since the onset of SCZ occurs typically in adolescence and early adulthood^47^, we examined 4 week (juvenile) and 12 week (young adult) mice. ∼9000 proteins were identified in the PSD fraction purified from cortex, including AKAP11 itself which was modestly (∼2x) enriched in the PSD fraction vs total lysate (Supplementary Fig. 4a,b). The number of differentially abundant proteins (DAPs, nominal *P* < 0.05) was greater in *Akap11^-/-^* than *Akap11^+/-^*mutants at 12 weeks but similar at 4 weeks, with approximately equal proportion of upregulated and downregulated DAPs overall (Fig. 4a, Supplementary Data 2). The DAPs were well correlated between *Akap11^+/-^*and *Akap11^-/-^* genotypes (Supplementary Fig. 4c) and weakly correlated between 4 and 12 weeks (Supplementary Fig. 4d).

**Figure 4:**
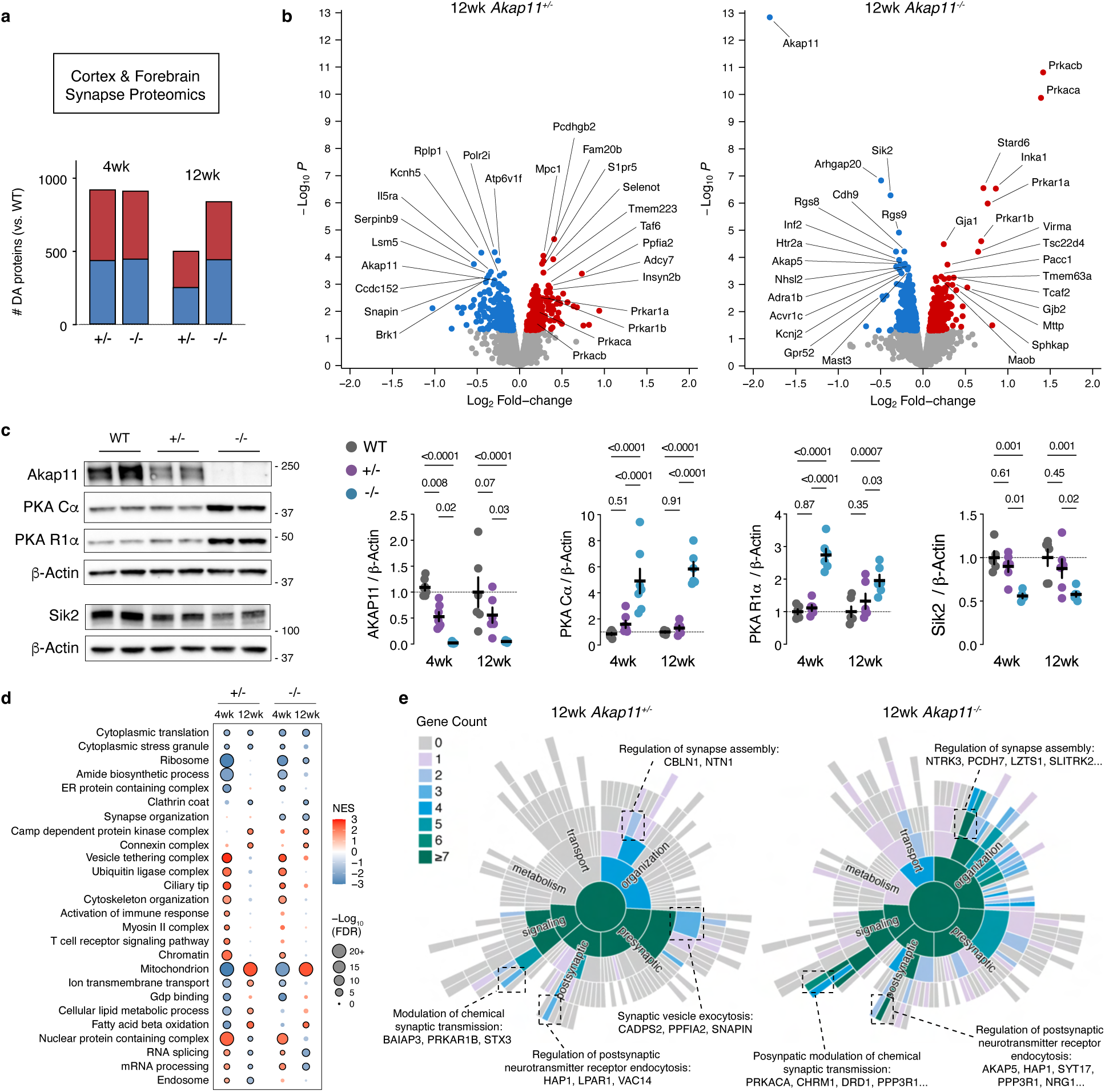
PKA accumulates in synapses of *Akap11* mutant mice. **a**, Number of DAPs across different ages and genotypes in synapse proteomics. Shading indicates upregulated (red) and downregulated (blue) DAPs, respectively. 4wk synapse proteomics data also appears in (Aryal et al., 2023)^50^. **b,** Volcano plots of 12wk synapse proteomics in *Akap11* mutants relative to WT. A moderated two-sample t-test was applied to the sample group datasets. Upregulated and downregulated DAPs are colored red and blue, respectively. **c,** Immunoblotting of synapse fractions with antibodies against AKAP11, PKA subunits, and SIK2. Right, quantifications for 4wk and 12wk animals. Data are represented as mean +/- SEM. Two-way ANOVA with Tukey’s post hoc test. See Supplementary Data 1 for statistics. **d,** A selection of GSEA analysis of synapse proteomics. Pathways displayed were significant at FDR < 0.05 in at least two age/ genotype conditions. **e,** Sunburst plots showing SynGO Biological Process pathways and the number of *P* < 0.05 proteins within these ontologies. Enrichment analysis is conducted with 1-sided Fisher’s exact test with all detected proteins as a background set^51^. For selected ontologies, representative DAPs in the ontology are shown. **b,d,e,g,** Two-way ANOVA with Tukey’s post hoc test. See Supplementary Data 1 for statistics.

In *Akap11^-/-^* synapses, the catalytic subunits PKA Cα and PKA Cβ were the two most upregulated proteins, at both 12 weeks (Fig. 4b) and 4 weeks, with PKA regulatory subunits upregulated to a lesser extent (Supplementary Fig. 4e,f). Immunoblotting confirmed strong accumulation of PKA Cα (∼5x) and R1α subunits (∼2-3x) in homozygous *Akap11^-/-^* PSD fractions (Fig. 4c) and modest elevation of PKA Cα (∼1.3x -1.6x) in heterozygous *Akap11^+/-^*synapses (Supplementary Fig. 4g). Besides PKA, many proteins showed changes in *Akap11^-/-^*synapse proteomics. Levels of SIK2, a PKA-regulated kinase involved in mood and circadian rhythms^48^ and ARHGAP20, a Rho GTPase-activating protein^49^, were significantly decreased in *Akap11^-/-^* synapses, at both 4 weeks and 12 weeks (Fig. 4b, Supplementary Fig. 4e,f). Notably, levels of AKAP11 itself were significantly decreased in *Akap11^+/-^* synapse fractions (Fig. 4b,c; Supplementary Fig. 4e), which is in contrast to the minimally altered levels of AKAP11 in *Akap11^+/-^* crude lysate (Fig. 3; Supplementary Fig. 3). This suggests that AKAP11 haploinsufficiency results in synapse-specific alterations to cellular signaling that are not apparent in bulk tissue readouts.

GSEA analysis of the synapse proteomic changes (Fig. 4d, Supplementary Data 3) revealed upregulation of “Vesicle tethering complex” and “Ubiquitin ligase complex” processes at 4 weeks (Fig. 4d, Supplementary Fig. 4h), which resembles the pathway changes observed in synapses purified from post mortem BD and SCZ human brain tissues^50^. These findings may relate to our observation that AKAP11 interacts with proteins involved in ER-Golgi transport and proteostasis (Fig. 2c,k). We further analyzed DAPs using gene ontology terms from SynGO^51^ to dissect the “sub-synaptic” processes that are altered in *Akap11* mutant synapses (Fig. 4e). DAPs in *Akap11^-/-^* mutant synapses were represented among pathways such as “Postsynaptic modulation of chemical transmission”, and “Postsynaptic neurotransmitter receptor endocytosis”. Relatedly, proteins in the “Clathrin coat” gene ontology were downregulated in 4wk and 12wk Akap11^+/-^ synapse proteomes (Supplementary Fig. 4i).

### Hyperphosphorylation of synapse proteins in *Akap11* mutant mice

Within *Akap11^-/-^* synapses, the levels of several protein kinases and phosphatases were altered, most prominently the accumulation of PKA catalytic subunits Cα and Cβ (Fig. 4b,c, Supplementary Fig. 4e). To quantify phosphorylation changes in synaptic proteins in an unbiased fashion, we analyzed PSD fractions purified from *Akap11* mutant cortex by quantitative phosphoproteomics. *Akap11* mutants showed a large number of differentially abundant phosphoproteins (DAPPs, nominal *P* < 0.05), which was higher in *Akap11^-/-^* than *Akap11^+/-^*, and higher at 4 weeks than 12 weeks (Fig. 5a, Supplementary Fig. 5a, Supplementary Data 2). We mapped changes in the synapse phosphoproteome against a reference of known and predicted kinase effectors^52^ to infer the kinases responsible. The vast majority of PKA phospho-substrates detected in the synapse phosphoproteomics data were increased in abundance in *Akap11^-/-^ ;* and a number of these phosphoproteins were also highly upregulated in *Akap11^+/-^* heterozygous mutant synapses (Fig. 5b). The sequences flanking the phosphosites showed strong enrichment of the basophilic RRXS*/T* motif, particularly in the upregulated DAPPs of *Akap11^-/-^* synaptic fractions at both 4 and 12 weeks (Fig. 5c), which matches the consensus amino acid sequence that is preferred for phosphorylation by PKA^53^. Together, the synapse proteomics and phosphoproteomics data show that PKA kinase activity as well as protein levels are abnormally elevated in the synapses of *Akap11* mutant mice.

**Figure 5:**
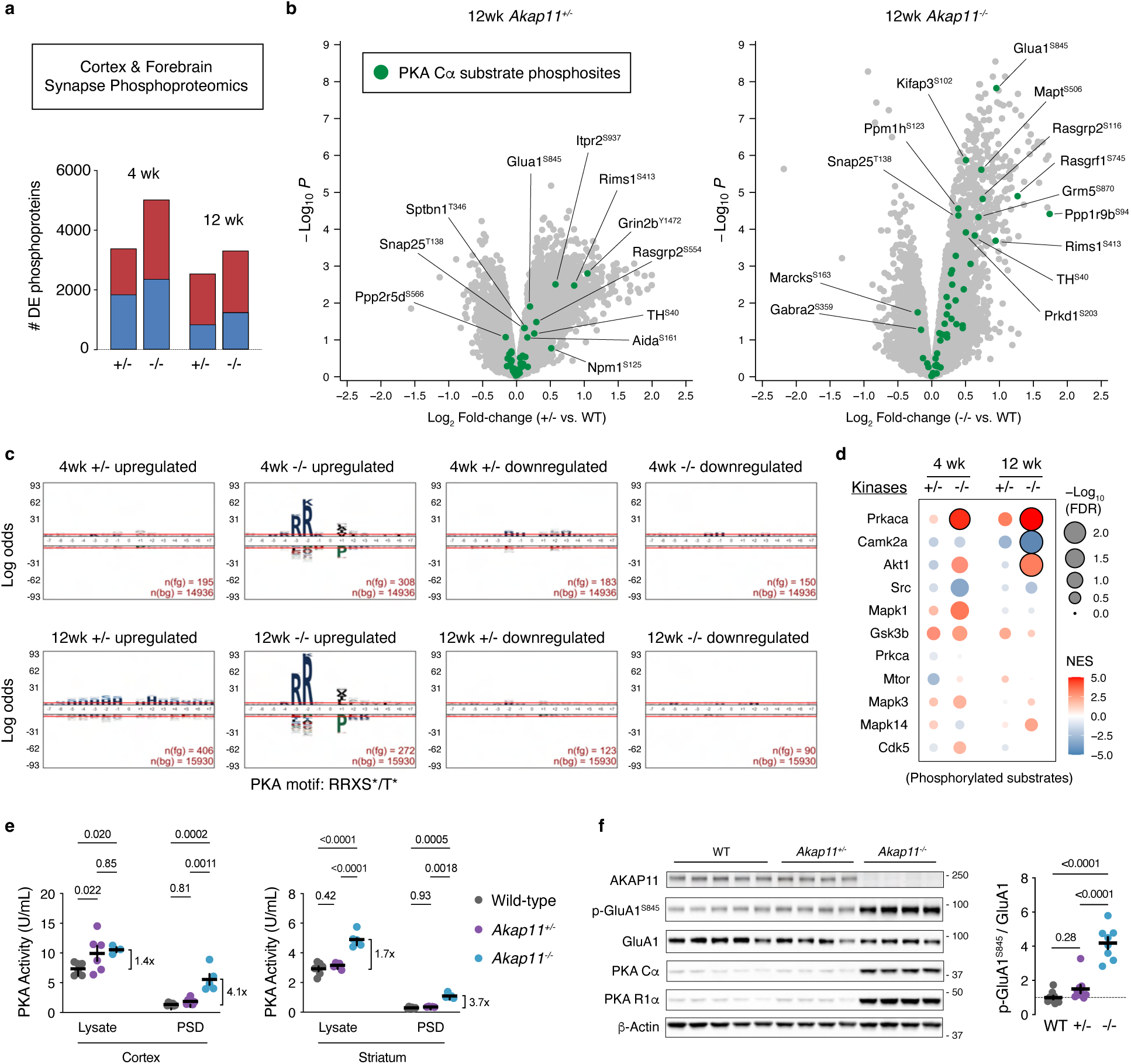
Phosphoproteomics of *Akap11* mutant synapses shows hyperphosphorylation of PKA substrates. **a,** Number of phosphoproteins reaching a significance threshold of nominal *P* < 0.05. Fractions of upregulated and downregulated DAPPs are highlighted in red and blue respectively. **b,** Volcano plots of 12wk synapse phosphoproteomics data for *Akap11^+/-^*and *Akap11^-/-^* vs. WT controls. Phosphoproteomic measurements are normalized to MS-proteomics measurements from the same samples. Known PKA Cα substrates from PhosphoSitePlus are labeled green. AKAP11 phosphopeptides are omitted from *Akap11^-/-^*plots, for clarity. **a, b,** A moderated two-sample t-test was applied to the datasets to compare WT, *Akap11^+/-^* and *Akap11^-/-^* sample groups. **c,** Motif analysis of peptides flanking the phosphorylation site of *P*<0.05 phosphosites in A. n(fg) are the number of foreground peptides, and n(bg) are the number of background peptides. Red lines indicate FDR significance, calculated by log odds enrichment. **d,** PTM-SEA of synapse phosphoproteomics data, using known kinase substrates from PhosphoSitePlus. Black borders indicate FDR significance. **e,** ELISA-based PKA activity measurements comparing total lysate and synapse fractions of cortex (left) and striatum-enriched tissue (right). One Unit (U) is defined by the manufacturer as the quantity of PKA that catalyzes the transfer of 1.0 pmol phosphate from ATP to substrate. Two-way ANOVA with Tukey’s post hoc test. **f,** Immunoblotting and quantification of AKAP11, PKA subunits, p-GluA1^S845^ and GluA1 in 12wk cortical total lysate. One-way ANOVA with Tukey’s post hoc test. **e,f,** Data are represented as mean +/- SEM. See Supplementary Data 1 for detailed statistical information.

Post-Translational Modification Signature Enrichment Analysis (PTM-SEA), which measures enrichment of kinase substrate sets (akin to GSEA using GO terms)^54^, also indicated elevation of phosphorylated PKA substrates in A*kap11^-/-^*, and to a lesser degree, *Akap11^+/-^* synapses (Fig. 5d). Phosphorylation of substrates by CaMK2A, a kinase involved in synaptic potentiation^55^ was significantly downregulated in *Akap11^-/-^* synapses at 12 weeks. Since AKAP11 is reported to inhibit GSK3β, it was intriguing to find that phosphorylation of *β*-Catenin (a well-known GSK3β substrate, encoded by *Ctnnb1*) was increased at Ser37 in *Akap11* mutants^56,57^ (Supplementary Fig. 5c). Phosphorylation of GSK3β itself on S9, a PKA phosphosite that inhibits GSK3β activity^58^, was significantly elevated in *Akap11* mutants (Supplementary Fig. 5f). Regulation of brain GSK3β by *Akap11* is especially relevant because lithium, an inhibitor of GSK3, is a mainstay treatment of BD^59^ and GSK3β overactivity has been implicated in SCZ animal models^60^.

To further test whether *Akap11* deficiency altered PKA activity in a compartment-specific fashion, we compared net PKA activity in total lysate versus synapse fractions, using a colorimetric enzyme-linked immunosorbent assay (ELISA) that measures phosphorylation of a PKA substrate. PKA activity was increased in both total lysates and synapse-enriched fractions of cortex and striatum (see Methods, Fig. 5e, Supplementary Fig. 5e). Notably, the relative increase of PKA activity is greater (∼3.9x) in synapse fractions than in total lysate (∼1.6x). Among the most highly changed phosphoproteins in *Akap11^-/-^* synapses was AMPA receptor subunit GluA1 (*Gria1*), which is phosphorylated by PKA at S845 (annotated as S863 in the UniProt mouse reference; Fig. 5b,f; Supplementary Fig. 5a,b). Phosphorylation of GluA1 at S845 is a well-known correlate of AMPA receptor trafficking to the synapse^61–65^. We confirmed by immunoblotting that GluA1^S845^ phosphorylation was sharply increased (∼4x) in brain lysates of *Akap11^-/-^*mutants, while total GluA1 protein showed little change (Fig. 5f, Supplementary Fig. 5g). Additional DAPPs in *Akap11^-/^*^-^ synapses that are known to be regulated by PKA include Snap25^T138^, a phosphosite required for presynaptic vesicular release^66^ and RIM1^S413^, phosphorylation of which is induced during synaptic potentiation^67–69^ (Fig. 5b). In *Akap11^+/-^*synapses, PKA substrate DAPPs included RIMS1^S413^ and ITPR2^S937^, a phosphosite that regulates intracellular calcium release^70^. The strength of synaptic transmission is regulated in large part by dynamic phosphorylation of synaptic proteins, including postsynaptic glutamate receptors subunits such as GluA1^71^. By mapping our phosphoproteomics data onto the phosphosites that are induced with chronic neuronal activation or silencing^72^, we found that many phosphosites associated with upscaling or downscaling of synaptic strength were also highly changed in *Akap11* mutants (Supplementary Fig. 5d), reflecting potential changes in synaptic function.

### Aberrant synaptic function in *Akap11* mutants

To directly examine synaptic function and plasticity in *Akap11* mutants, we performed electrophysiology recordings from acute hippocampal slices (6-8 week old mice). Under basal conditions, the stimulation input/output relationships (stimulation of Schaffer collaterals to CA1) of *Akap11^-/-^* mice were indistinguishable from WT (Supplementary Fig. 6a). The amplitude of early long-term potentiation (LTP) induced by theta burst stimulation (TBS) was modestly but significantly reduced in *Akap11^-/-^* mice (Supplementary Fig. 6b). We speculate this phenotype could be due to partial occlusion of LTP by hyperphosphorylated GluA1 at S845^64,71,73^*. Akap11^-/-^*slices also showed heightened paired-pulse facilitation (Supplementary Fig. 6c), suggesting reduced presynaptic release probability, which might relate to altered RIM1^S413^ and Snap25^T138^ phosphorylation seen in the synapse phosphoproteomics (Fig. 5b). The electrophysiologic phenotype in heterozygous *Akap11^+/-^* hippocampus was generally not significantly different from WT, though there was a trend toward reduced magnitude of early LTP (Supplementary Fig. 6b).

### Altered synaptic genes and striatal circuits revealed by transcriptomic analysis

*Akap11* mRNA is broadly expressed across the brain and across postnatal development in wild-type mice (Supplementary Fig. 7a). To query the brain regions/circuits and molecular pathways that are impacted by *Akap11* loss of function, we conducted bulk RNA-seq on PFC, SSC, Str, Thal, Hipp, SN, as well as single-nucleus RNA sequencing (snRNA-seq) on PFC and Str, at 4 weeks and 12 weeks (Fig. 6a, Supplementary Fig. 7b,c). By bulk RNA-seq, we found more differentially expressed genes (DEGs; defined as genes that changed in either direction with FDR <0.05, Supplementary Data 4) in *Akap11^-/-^* than *Akap11^+/-^*, and at 4 weeks than at 12 weeks of age (Fig. 6b). Based on DEG count, the most impacted regions in *Akap11^-/-^* were striatum, thalamus and PFC, whereas thalamus was the most affected brain region in *Akap11^+/-^*. The striatum had by far the most DEGs, at either age, in *Akap11^-/-^*. Only a few DEGs were common across brain regions (Supplementary Fig. 7d), and the correlation of Log_2_-Fold-Changes of nominally significant genes was low between different brain regions, and modest between 4 and 12 weeks for the same brain region (Supplementary Fig. 7e). Thus the transcriptional consequence of *Akap11* mutation varies with the brain region and developmental context. Nonetheless, GSEA of bulk transcriptomic changes revealed consistent dysregulation of biological pathways related to synapses, mitochondria/oxidative phosphorylation, ribosomes, and RNA processing/splicing, which was particularly evident in 4wk *Akap11^+/-^* (Fig. 6c, Supplementary Data 5). The “Synapse” GO term was consistently downregulated in PFC and SN at both 4 and 12 weeks in *Akap11^+/-^* and *Akap11^-/-^* mutants (Fig. 6c). Immune-related pathways were upregulated in the 4wk PFC and SN, which is notable in the context of synaptic pruning by glia during development and disease^74^ and SCZ associations with immune-related genetic loci^75,76^. Two pathways required for cellular energetics and upkeep, “Oxidative phosphorylation” (Ox-phos) and “Ribosome”, were changed in correlated fashion across brain regions, similar to observations in another SCZ rare variant mouse model *(GRIN2A)*^12^; these pathways are also altered in human SCZ postmortem samples^77,78^. The “ribosome” and “oxidative phosphorylation” pathway (a mitochondrial process that generates ATP) were down-regulated at 4 weeks in the PFC of both *Akap11^+/-^ and Akap11^-/-^*mutants (Fig. 6c). We investigated changes to neuronal activity at the brain region level in *Akap11* mutant mice. As a surrogate measure of neuronal activity, we conducted GSEA of the bulk RNA-seq data using a set of activity-regulated genes (ARGs, Fig. 6c)^12,79^. At 4 weeks, ARGs were upregulated in the striatum of *Akap11^+/-^* and downregulated in the SSC and PFC of A*kap11^-/-^* (though the latter did not reach significance). The consistent downregulation of the “Synapse”, “Ox-phos” and “ARGs” GO terms in PFC is suggestive of hypofunction of PFC that is reported in fMRI studies of humans with SCZ and BD^80,81^.

**Figure 6:**
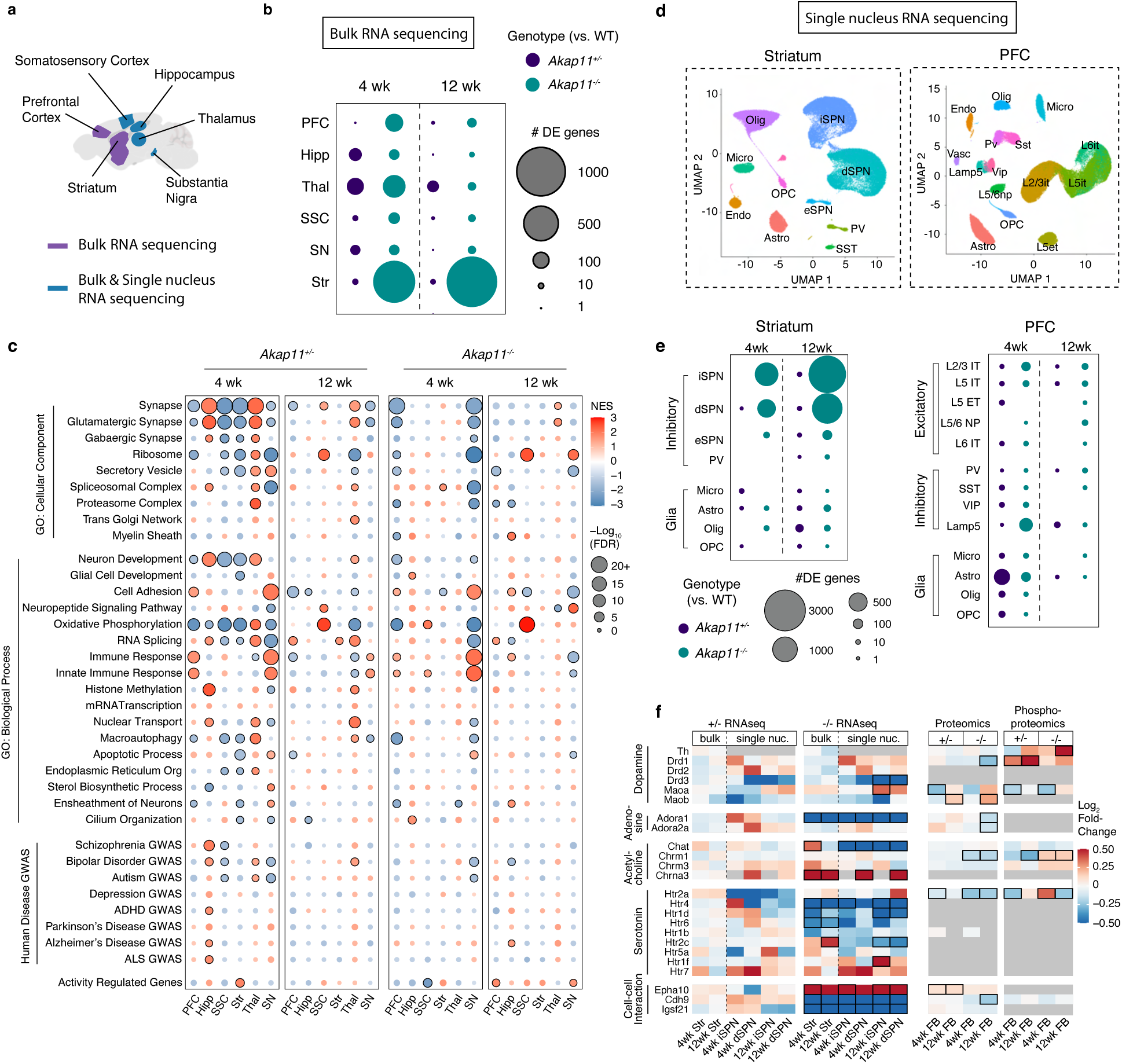
Transcriptome-wide impact of *Akap11* deficiency across ages, brain regions and cell types. **a,** Schematic of brain regions collected for RNA sequencing. **b,** Number of differentially expressed genes (DEGs) in bulk RNA sequencing across brain regions and ages. Experiments are n= 4 to 6 per condition (see Methods). Area of circles represents DEG counts, defined by FDR adjusted *P*-values < 0.05, for *Akap11* mutants compared to WT controls. Two-tailed Wald test, with B-H FDR correction. **c,** GSEA of bulk RNA sequencing, showing MsigDB pathways of interest, genes associated with psychiatric and neurological conditions by GWAS^12,82^, and a curated set of activity-regulated genes^79^. Black outline indicates FDR < 0.05 significance. GSEA uses a Kolmogorov-Smirnov test, two tailed, with B-H FDR correction applied. **d,** Schematic of snRNA-seq experiments in PFC and striatum, with UMAP plots of major cell types identified at 12w by snRNA-seq. Experiments are n= 4 per condition, with the exception of 12w Str, with n=5 WT. **e,** Number of DEGs in snRNA-seq experiments, defined by having an FDR adjusted *P*-value < 0.05. Comparisons were calculated using two-tailed likelihood ratio test, which were FDR corrected with B-H adjustment. **f,** Heatmap showing Log_2_FC values for a selection of neuromodulation-related genes, plotted across transcriptomics, proteomics, and phosphoproteomics measurements. Genes indicated in the phosphoproteomics columns reflect the most significantly changed phosphosite per gene. *P* < 0.05 proteins and FDR < 0.05 transcripts are outlined with a black box.

We queried the bulk RNA-seq changes in *Akap11* mutants for enrichment of SCZ- and BD-associated risk genes identified from human genome-wide association studies (GWAS) (Fig. 6c). The most significant enrichments occurred for BD risk genes, and interestingly, risk genes associated with multiple psychiatric disorders were most enriched in *Akap11^+/-^* heterozygous mutants at 4 weeks of age (Fig. 6c). Though reaching lower statistical significance across fewer brain regions, the pattern of SCZ risk gene enrichment had an overall similar pattern to BD. Besides SCZ and BD, we also observed similar GSEA signals for genes associated with ASD and neurodevelopmental disorders^37,82–84^. Overall, these results support that *Akap11* functionally engages with other SCZ and BD and neurodevelopmental risk genes, and are consistent with the known genetic overlap between SCZ, BD, and neurodevelopmental disorders (NDD)^5,85^.

### Major disturbances to striatal neurons revealed by single nucleus RNA-seq

In WT mice, *Akap11* mRNA is expressed in excitatory and inhibitory neurons in all brain regions, as well as in glia (Supplementary Fig. 8a). To investigate the cell type-specific effects of *Akap11* deficiency, we conducted single-nucleus RNA sequencing (snRNA-Seq) in the PFC and striatum at 4 and 12 weeks (Fig. 6d, Supplementary Fig. 8b, Supplementary Data 6). There were no major changes in cell type proportions in *Akap11* mutant PFC or striatum (Supplementary Fig. 8b). Although all cell types in PFC and striatum showed transcriptomic changes in *Akap11* mutants, the largest numbers of DEGs (FDR < 0.05) were found in the major neuron-types of the striatum of *Akap11^-/-^,* namely the inhibitory direct- and indirect-pathway spiny projection neurons (dSPNs, iSPNs; Fig. 6e). Transcriptomic alterations in *Akap11^-/-^* mutants were highly correlated between iSPNs and dSPNs (Supplementary Fig. 8c,d) and between 4 and 12 weeks of age (Supplementary Fig. 8d,e). Consistent with bulk RNA-seq findings (Fig. 6b), *Akap11^+/-^* had much less effect than *Akap11^-/-^*on SPNs in snRNA-seq (Fig. 6e, Supplementary Fig. 8d,f). Nearly all PFC neuron subtypes showed downregulation of the “Synapse” GO term (Supplementary Fig. 9a, Supplementary Data 7), which aligns with bulk RNA-seq (Fig. 6c). Unlike striatal neurons, which are much more impacted by homozygous than heterozygous *Akap11* mutation, glia in the PFC and striatum (astrocytes, oligodendrocytes and microglia) showed comparable numbers of DEGs genes between *Akap11^+/-^* and *Akap11^-/-^* (Fig. 6e). *Akap11* mutant astrocytes showed downregulation of the “Sterol Biosynthesis” pathway, particularly in the 4 week *Akap11^+/-^* PFC and *Akap11^-/-^*striatum (Supplementary Fig. 9b). Across regions, ages and genotypes, decreases in synapse-related gene expression in neurons often occurred alongside decreased expression of cholesterol biosynthesis genes in astrocytes (Supplementary Fig. 9c), suggesting coordinately disturbed neuron-glia gene expression, as has been observed in postmortem human SCZ brain tissue^86^.

The largest changes in the *Akap11* mutant SPNs highlighted several neuromodulatory systems, such as serotonin (example genes: *Htr2a*, *Htr4*), adenosine (*Adora1*), acetylcholine (*Chat*, *Chrna3*), and dopamine (*Cartpt*) (Fig. 6f; Supplementary Fig. 10a,b). Intriguingly, the serotonin receptor HTR2A, which is a target of hallucinogenic drugs^87,88^, was downregulated in both *Akap11^+/-^*synapse proteomics and snRNA-seq of striatum SPNs, and *Htr4* was robustly downregulated in bulk RNA-seq of striatum and in snRNA-seq of striatal SPNs (Fig. 6f; Supplementary Fig. 10a,b). Adenosine receptor ADORA1, a G-protein coupled receptor that inhibits cAMP production, was downregulated in *Akap11^-/-^* synapse proteomics as well as in bulk RNA-seq and snRNA-seq data in striatum and SPNs. Choline acetyltransferase (*Chat*) was also prominently downregulated in the striatum of *Akap11^-/-^* (Fig. 6f). *Chat* participates in the synthesis of acetylcholine, and signaling through muscarinic acetylcholine receptors is implicated in SCZ pathophysiology^89,90^. Transcellular signaling factors *Epha10*, *Cdh9*, and *Igsf21* were among the most altered genes in the *Akap11^-/-^*striatum (Fig. 6f). *Epha10* is a receptor tyrosine kinase that acts as an axon guidance molecule^91,92^. *Cdh9,* which decreased consistently across proteomics and transcriptomics measurements of *Akap11^-/-^*, is involved in calcium dependent cell-cell adhesion^93^. Together, these alterations in neuromodulation and trans-cellular signaling indicate potentially aberrant cell-to-cell interactions in the striatum of *Akap11^-/-^*mice.

Dopamine is a key neuromodulator in the striatum^94^. Excessive or aberrant dopaminergic signaling in the striatum is a long-standing hypothesis in pathophysiology of schizophrenia and other psychiatric disorders, underpinned by the antipsychotic efficacy of D2 dopamine receptor antagonists^95^ .We found several dopamine-related genes, proteins, and posttranslational modifications that were altered in *Akap11* mutants (Fig. 6f), including: (i) increased phosphorylation of tyrosine hydroxylase (TH) at S40, which positively regulates its dopamine biosynthesis activity^96,97^; (ii) reduction of protein and increased phosphorylation of dopamine receptor DRD1; (iii) increased phosphorylation of RASGRP2^S116^ in 12 week *Akap11^-/-^* synapses (Fig. 5b), which is consistent with enhanced DRD1 signaling^98^; (iv) elevated MAO-B protein in synapse fractions, an enzyme that degrades dopamine, v) increased proportion of *Drd2-*expressing (Drd2^+^) neurons in *Akap11^+/-^* relative to wild-type at 4 and 12 weeks (Supplementary Fig. 10c). One dopamine-regulated transcript, *Cart* (Cocaine- and amphetamine-regulated transcript)^99,100^ was highly elevated in the *Akap11^-/-^* striatum (bulk RNA-seq) and in striatal SPNs and PFC excitatory neurons (snRNA-seq) (Supplementary Fig. 10d) which may reflect elevated dopamine signaling in striatum and in PFC of *Akap11* mutants.

### *In vivo* striatal PKA dynamics and dopamine signaling are regulated by AKAP11

Two of the most striking phenotypes of *Akap11* mutant mice are highly elevated PKA protein (Figs. 3-4) and strong transcriptomic changes in the striatum and SPNs (Fig. 6). PKA is a central signaling molecule in the striatum, acting in the signal transduction pathway downstream of dopamine receptors^94^. To assess real-time dynamics of PKA activity in live animals, we used fluorescent lifetime fiber photometry measurements of FLIM-AKAR, a fluorescence lifetime reporter of net PKA activity^101,102^ that was virally expressed in the nucleus accumbens (NAc; within the ventral striatum) (Fig. 7a, Supplementary Fig. 11a-c). In awake, behaving *Akap11^-/-^*mice, PKA activity was elevated across 10 minute recording sessions in an open field arena, (Fig. 7b) which was consistent across weeks of measurement (Supplementary Fig. 11d). As a negative control, there were no differences across *Akap11* genotypes in the lifetime of FLIM-AKAR^T391A^, a mutated variant of that is insensitive to PKA activity (Fig. 7c).

**Figure 7:**
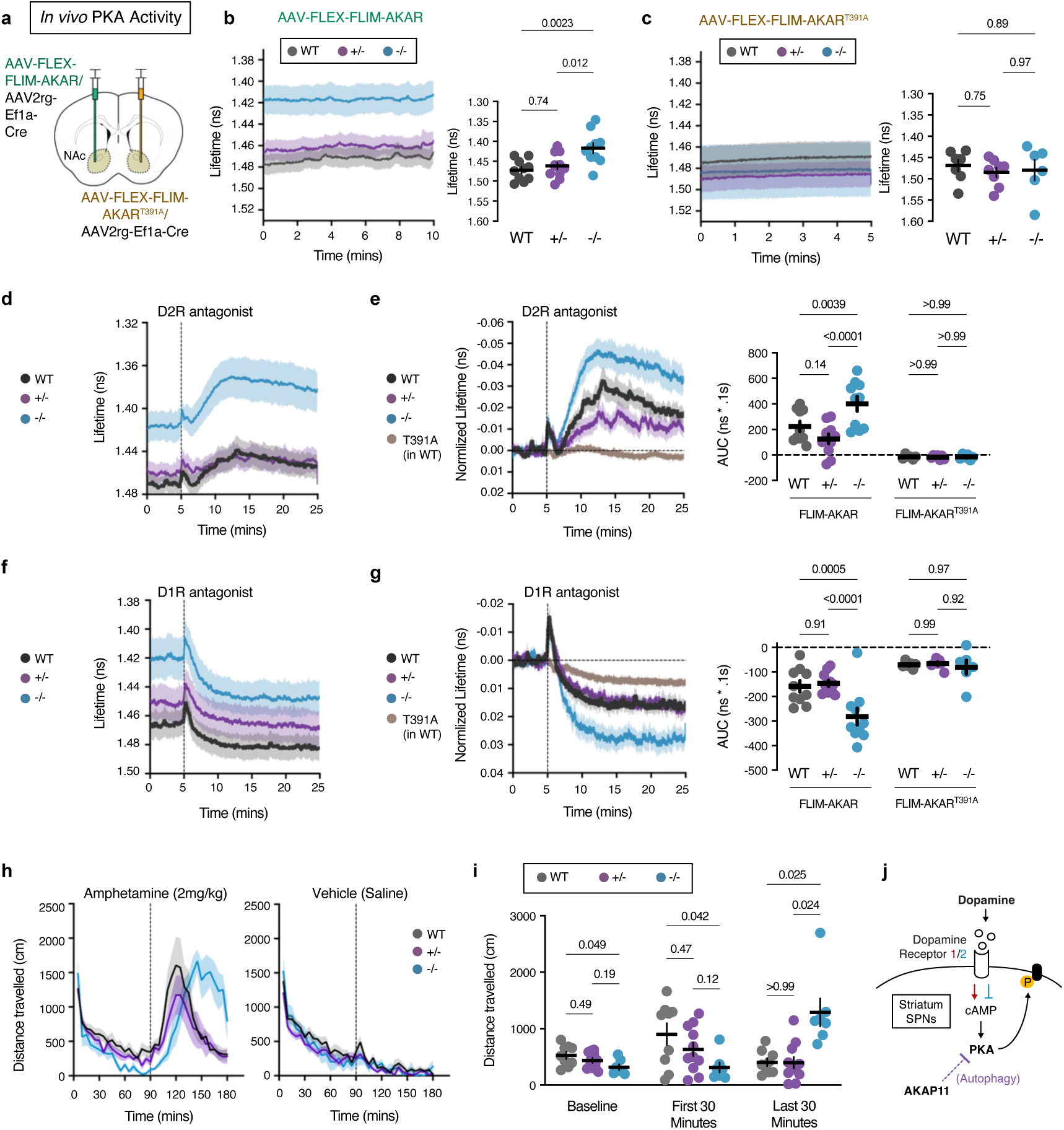
Fluorescence lifetime photometry (FLiP) in *Akap11* mutant mice reveals constitutively elevated PKA activity and functional alterations to striatal circuitry. **a,** Schematic overview of the viral injection strategy. **b,** Fluorescence lifetime of FLIM-AKAR in WT or *Akap11* mutant mice. The y-axis is inverted to reflect that decreased fluorescence lifetime corresponds to increased net PKA activity. **c,** Fluorescence lifetime of FLIM-AKAR^T391A^ in WT or *Akap11* mutant mice. **d,** Fluorescence lifetime changes of FLIM-AKAR in response to D2R antagonist. **e,** Fluorescence lifetime changes of FLIM-AKAR in response to D2R antagonist, normalized to each genotype’s baseline level of activity. **f,** Fluorescence lifetime changes of FLIM-AKAR in response to D1R antagonist. **g,** Fluorescence lifetime changes of FLIM-AKAR in response to D1R antagonist, normalized to each genotype’s baseline level of activity. **h,** Distance traveled after administration of amphetamine (2 mg/kg) or saline to WT or *Akap11* mutants. **i,** Quantification of distance traveled during baseline (0-90mins), 90-120mins, and 150-180mins. Repeated measures two-way ANOVA with Tukey’s post hoc test. **j,** Schematic of dopamine to PKA regulation by *Akap11*. **i,** Data are represented as mean +/- SEM, with the shaded regions or error bars representing SEM.

Dopamine receptors are enriched in the striatum, and excessive dopamine signaling in striatum may drive positive symptoms (e.g. hallucinations) of SCZ^103–105^, which is supported by the clinical efficacy of dopamine receptor 2 (*Drd2*) antagonists for positive symptoms^95^. To evaluate how PKA is dynamically regulated by dopamine in *Akap11* mutant mice, we administered dopamine receptor antagonists via intraperitoneal injection. Dopamine acting on D2-type receptors inhibits cAMP production and thereby inhibits PKA^94,106^. As expected, net PKA activity was increased in striatal neurons following administration of D2 antagonist (eticlopride hydrochloride, 0.5 mg kg^−1^) (Fig. 7d). Despite the higher baseline activity of PKA activity in *Akap11^-/-^*striatum, phosphorylation of the FLIM-AKAR sensor was further enhanced by D2 antagonist in *Akap11^-/-^* mutants (Fig. 7e). Dopamine acting on D1-type receptors stimulates cAMP synthesis and activates PKA^94,106^. As expected, delivery of D1 antagonist (SKF83566, 3 mg kg^−1^) suppressed net PKA activity in striatum in all 3 genotypes (Fig. 7f). When normalized to the pre-drug baseline level, D1R antagonism caused a more pronounced inhibition of PKA activity in *Akap11^-/-^* mice (Fig. 7g). Thus loss of *Akap11* results in markedly higher basal PKA activity; however, this increased basal activity does not occlude further activation or block inhibition of PKA by dopamine receptor drugs, but rather expands the dynamic range of PKA activation or inhibition by endogenous dopamine.

Dopamine in the striatum regulates multiple aspects of behavior, including movement, reward processing, cognition and mood^107^. Dopamine levels induced by amphetamine are higher in the striatum of schizophrenic patients^108^. We tested locomotor response to subcutaneous amphetamine administration, which depends on dopamine and is often used as a behavioral measure of dopamine sensitivity in rodents (Fig. 7h,i). Wild-type mice, as expected, showed amphetamine-induced hyperactivity (AIH; measured by distance traveled). During the first 30 minutes after amphetamine administration, *Akap11^-/-^* mice displayed less AIH than WT littermates; however, *Akap11^-/-^* mice showed a much more sustained hyperactivity response, such that at 60-90 minutes after amphetamine administration, *Akap11^-/-^* mice still showed high levels of locomotion, when WT animals had returned to baseline (Fig. 7h). During this prolonged period of hyperactivity, *Akap11^-/-^* mice spent more time in the center of the open field, and exhibited more rearing movements than WT or heterozygous mutants (Supplementary Fig. 12 a-f). In conjunction with FLIM-AKAR and behavioral readouts, these *in vivo* pharmacological experiments suggest that the strength and temporal dynamics of dopamine-to-PKA signaling in striatal SPNs are abnormally high and excessively prolonged in *Akap11* mutants (Fig. 7j).

## Discussion

Genetic variation contributes strongly towards schizophrenia and bipolar disorder risk. Psychiatric symptoms ultimately emerge from dysfunction of specific neural circuits; however the mechanisms by which genetic variants lead to molecular-cellular disturbances and then to circuit dysfunction remain unclear. In this study, we examined mice with mutation in *Akap11*, a risk gene for schizophrenia and bipolar disorder. Our unbiased multi-omic studies revealed that AKAP11 loss of function results in a remarkable elevation of PKA activity in the brain, especially at synapses; and the striatum, the major site of dopaminergic signaling and integration in the brain, is particularly impacted by *Akap11* deficiency.

Hyperdopaminergic signaling in striatum is strongly implicated in the positive symptoms of psychotic disorders. Our study focuses attention on factors downstream of dopamine and dopamine receptors - namely, PKA dysregulation in SPNs --as an important mechanism by which striatal dopaminergic signaling can be exaggerated. Since PKA is elevated across the brain in *Akap11* mutants, we expect that excessive PKA activity also contributes to pathophysiological disturbance in other brain regions and circuits beyond the striatum - such as the PFC and thalamus, where we also saw major transcriptomic changes.

It is important to note that, in comparison with homozygous *Akap11* knockout, the phenotypes of *Akap11* heterozygous mutants are relatively subtle. With the exception of transcriptomics changes in thalamus and in glial cell types (astrocytes, oligodendrocytes and microglia), we observed much larger molecular, physiological and behavioral phenotypes in homozygous than in heterozygous mutants. However, the phenotypes that were most prominent in *Akap11^-/-^*, such as elevated PKA, were also seen in *Akap11^+/-^*, albeit to a lesser degree. The modest phenotype in *Akap11* heterozygotes could be explained by the fact that *Akap11* haploinsufficiency resulted in only slight changes in AKAP11 protein levels. This latter observation suggests that there are proteostatic mechanisms operating in heterozygous mutants to buffer against sharper losses of AKAP11 protein and function. Since *Akap11* regulates autophagy, it would be especially interesting to test whether cellular stressors, which can stimulate autophagy, could exacerbate the reduction of AKAP11 protein and amplify the downstream consequences such as elevation of PKA activity in *Akap11* heterozygous mutants, thereby offering a potential way for environmental triggers to induce a stronger disease phenotype.

Across transcriptomic measurements in varying brain regions and ages, the most consistent alterations were among genes related to oxidative phosphorylation, mitochondrial respiration, ribosomes and RNA processing / splicing. Intriguingly, selective autophagy of PKA regulatory subunit R1α through interactions with AKAP11 increases PKA activity and mitochondrial metabolism in the context of glucose starvation^23^. We have also uncovered direct evidence for abnormal lipid metabolism in the *Akap11* mutant brain^109^. Future work should explore whether *Akap11* mutant brains respond differently to environmental, metabolic and cellular stressors.

There are likely to be hundreds of genetic variants underlying SCZ and BD. Because these genes are diverse in molecular function, their most proximal molecular mechanism would be expected to be varied and perhaps unique to each genetic variant. Thus, it is essential to identify downstream convergent cellular and circuit disturbances that are shared among different SCZ and BD risk genes. Our findings highlight one mechanistic pathway by which a specific genetic variation leads to neural circuit dysfunction. In this case, loss of function of *Akap11*, which is required for autophagy-degradation of PKA^23,45^, leads to elevated PKA kinase activity in neurons, resulting in unbalanced dopamine-to-PKA signaling in SPNs of striatum. Another mouse model of a rare variant SCZ gene (*GRIN2A*)^5^ also shows evidence of hyperdopaminergic signaling in striatum, but via a mechanism different than exaggerated PKA activation^12^. In *SETD1A* mutant cell lines and *Trio* mutant mice, which are also genetic models for schizophrenia^5^, there is elevated PKA activity that results in synaptic dysfunction^13,110^. Further, the cAMP synthesizing enzyme adenylyl cyclase 2 (*Adcy2*) is a significant genetic association for bipolar disorder^8^. Thus, human genetics and mechanistic studies corroborate a plausible role of PKA dysregulation in psychiatric disorders.

D2 receptor antagonists are efficacious for treatment of SCZ positive symptoms; this seems counterintuitive to the results of our fiber photometry experiments, in which D2 antagonism further exacerbated the PKA hyperactivity phenotype of *Akap11^-/-^*. Although used in the clinic for decades, the physiologic mechanism for the efficacy of D2 antagonists in SCZ is still unclear^111,112^. Clinically effective antipsychotics (including D2 receptor antagonists) better correlate with the suppression of D1-expressing neuron (dSPN) activity, rather than the suppression of D2-expressing neurons (iSPNs)^113^, suggesting that the activity of D1-expressing neurons is more relevant to schizophrenia pathophysiology. Since dSPN neuron activity is enhanced by PKA activation^98^, it is reasonable to predict that dSPN activity is increased in *Akap11* mutants. Future work should directly measure neuronal activity in D1 and D2 SPNs, and how their activity levels are affected by antipsychotic medicines and mood-stabilizing drugs like lithium.

A separate study of *Akap11* mutant mice also found elevated PKA activity in synapse-enriched fractions, but decreased PKA activity in cytosolic fractions of the brain^114^. This apparent difference from our results could be accounted for methodological differences, such as full body germline knockout of *Akap11* (this study) versus neuron-specific knockout in Lee et al. (2025) Another possibility is that PKA activity is differently regulated by AKAP11 across cellular compartments, which should be the subject of future investigation.

Our study, which started with unbiased -omics approaches, highlights the multi-faceted molecular control of dopaminergic signaling in striatum – already much implicated in psychosis pathophysiology by human and mouse model studies – as a worthy focus of future investigation. Our multi-omic analysis of the *Akap11* mouse model, together with similar investigation of other high penetrance SCZ and/or BD genes, should enable additional hypothesis-driven experiments and lead to a more nuanced understanding of the complex and heterogeneous mechanisms underlying psychiatric disease.

## Methods

### Animals

Experiments were approved by the Broad Institute IACUC (Institutional Animal Care and Use Committee) and conducted in accordance with the NIH Guide for the Care and Use of Laboratory Animals (0259-10-19-1, 0288-11-20-1 and 0008-06-14). Heterozygous *Akap11* mice (B6.Cg-Akap11[tm1.2Jsco/J]) were obtained from The Jackson Laboratory following cryorecovery (strain #028922). As previously described^30^, these mice have deletions of exons 6 and 7 of the mouse *Akap11* gene to create a knockout allele. *Akap11* mutant mice were deposited by the donor lab after 6 generations of backcrossing to the C57BL/6J background. We confirmed the genetic background of the animals to be greater than 90% C57BL/6J through SNP analysis (MiniMuga panel, Transnetyx). To establish the colony, founders were crossed to WT C57/BL6J mice (Jackson Laboratory, #000664) for one generation. Subsequently, heterozygous mice were crossed with each other to produce wild-type (WT, +/+), heterozygous (Het, +/-) and knockout (KO, -/-) littermates used for experiments. Mice were housed at AAALAC-approved facilities on a 12-hour light/dark cycle, between 30-60% humidity, 67°-73° Fahrenheit, with food and water available ad libitum. For fluorescence lifetime photometry experiments (FLiP), mice were housed in a 12h/12h light/dark reverse cycle. FLiP and pharmacological experiments were conducted in compliance with protocols approved by the Harvard Standing Committee on Animal Care (IS-00000571-6), adhering to guidelines outlined in the US National Institutes of Health Guide for the Care and Use of Laboratory Animals.

### Homecage locomotor activity monitoring

Male and female mice aged P23-P34 were transferred to Digital Ventilated Cages (DVC, Tecniplast) and were single housed throughout the duration of the experiment. Briefly, DVC racks use positional sensors beneath each cage that can detect disturbances that correlate with animal movement^25,115^. The DVC Analytics web platform was used to generate animal locomotor indices, which calculate the percentage of 12 electrodes that measured movement over a 1-minute bin. For the locomotor activity index, a value of 1 means 12 electrodes measured movement, and a value greater than 1 means that there were > 12 detected movements. The experiment included n=11 WT males, n=12 WT females, n=12 *Akap11^+/-^* males, n=14 *Akap11^+/-^*females, n=10 *Akap11^-/-^* males, and n=13 *Akap11^-/-^* females. For a subset of animals, a running wheel was introduced for 1-2 weeks during the homecage study. Running wheel usage was assessed during the first 6 days of introduction. For statistical comparisons of locomotor activity index (Fig. 1b,c), locomotor activity index data was excluded on cage change days, and days where the running wheel was present.

### Motion Sequencing (MoSeq) behavioral recording

Mice were tested in a Motion sequencing (MoSeq) open field assay as previously described^12^. Briefly, a matte black bucket with a Microsoft Kinect Camera attached overhead was used to perform the recordings. Two days prior to the recording session, mice were habituated for 10 minutes in the recording arena. On the day of the recording session, all mice being recorded that day were placed in the room 30 minutes prior to the first recording. Each mouse was individually recorded for 30 minutes. Prior to the first recording, the bucket was wiped with 70% ethanol and allowed to dry for 5 minutes. This was repeated after each subsequent mouse. Between mice, the arena was cleaned and the sides of the bucket were sanded down with sandpaper as to limit the glare leading to background noise during analysis. Mice were tested after the lights-off time (Zeitgeber Time 12, ZT12) since mice were nocturnal. All tests occurred between ZT12-ZT18. Mice were 12 weeks (+/- 1 week) of age. Sample sizes were 11 WT, 11 *Akap11^+/-^*and 10 *Akap11^-/-^* mice. Moseq syllable videos were manually annotated by five independent observers as “Grooming”, “Rearing”, “Pausing”, or “Turning, lunging or Darting”. Moseq data has been uploaded to a public repository (https://zenodo.org/records/17094753).

### MoSeq Computational Analysis

MoSeq is a tool that employs an unsupervised machine learning algorithm to characterize and quantify complex mouse behavior. It breaks down spontaneous mouse behavior into sub-second, regulated, and fine-tuned behavioral motifs called syllables^28^. Computational analysis of recordings as generated above were analyzed using the Depth MoSeq pipeline available here: http://www.moseq4all.org/. Extraction was performed with default configuration parameters and minimum mouse height set to 12 to reduce visual glare in the recordings. After generating a robust model with selected kappa value (kappa=599484, trained with 1000 iterations of Gibbs sampling) as outlined in the pipeline, 51 syllables were produced, which explained 99% of the total frames. These syllables were manually labelled into four larger groups of mouse behavior: pausing, grooming, rearing/climbing, and darting/lunging/turning. After assigning each frame of each video a syllable label, the number of occurrences of each syllable (syllable usage) and the number of transitions from one syllable to another were quantified. Syllables were then ordered and labelled by the fraction of times they were used across all sessions, as seen in Fig. 1e and 1f, with syllable 0 being the most used syllable and syllable 51 being the least used behavioral motif.

### Discriminatory fear conditioning

Mice were habituated to a sound attenuated conditioning chamber, “Context A,” 30.5 × 24.1 × 21.0 cm^3^ dimensions with an automated freezing analysis software (Med Associates) and were presented with 10X alternating trials of 20 second 3 KHz and 10 KHz auditory tones (75 dB sound pressure level). The next day, mice underwent fear conditioning in the same chamber with the same tones. Mice were exposed to 10X trials of stimulus–unconditioned stimulus pairings (tone-foot shock) with the 3 KHz tone or 10 KHz tone serving as the reinforced conditioned stimulus and the other tone serving as the neutral non-reinforced stimulus (counterbalanced). The aversive unconditioned stimulus was a 0.3 mA foot shock co-terminating with the last 1.5 seconds of the CS+. Context A was cleaned before and after behavioral sessions with chlorine dioxide sterilant solution (Clidox-S). Memory recall and extinction sessions occurred 24 and 48 hours. later in a novel “Context B” consisting of the same chamber with white contextual inserts (white curved wall and white floors) and vanilla extract to serve as a distinct olfactory cue. Mice were presented with 15X trials of non-reinforced CS+ and CS- tones per session. Context B was cleaned with 70% ethanol before and after each session. All CS+ and CS- tones were presented in an alternating fashion with a 60 second intertrial interval. Memory recall and discrimination to the CS+ and CS- tones was scored as average freezing behavior (cessation of all movement other than respiration for >1 second) during the first 5X tones presentations during the first recall session. To compare sensory discrimination normalized to each animals’ behavioral performance and overall fear level we generated a discrimination index using the percent time freezing, defined as: (CS+ freeze - CS- freeze) / (CS+ freeze + CS- freeze).

### Behavioral Battery

Fifty-five AKAP11 mice (n=19 WT, n=22 *Akap11^+/-^*, n=14 *Akap11^-/-^*) aged between 12-16 weeks were used in a battery of behavioral assays including sucrose preference test, forced swim test, and prepulse inhibition. Animals were grouped housed in a maximum of 5 animals per cage, under normal lighting (lights on 7am-7pm) and ad libitum supply of food and water. All procedures were approved by the Institutional Animal Care and Use Committee at Boston Children’s Hospital and were in accordance with the National Institutes of Health Guide for the Care and Use of Laboratory Animals.

### Sucrose preference test

For the sucrose preference test: on day 1, mice were individually housed and were habituated to 2% sucrose for 24 hours. All bottles were pre-weighed and each mouse got two bottles filled with 2% sucrose solution. The bottles are retrieved and weighed at the end of the habituation day. On day 2 and 3, mice were individually housed and underwent a sucrose preference test.

One of the bottles was filled with 2% sucrose solution while the other was filled with water. The position of the sucrose/water bottle was reversed at the end of day 2, and the bottles were weighed at the end of each day to determine water/sucrose consumption. Sucrose preference test was conducted once at the beginning of the behavioral test battery and was repeated once after other behavioral assays (forced swim test and prepulse inhibition) were completed. The experimenter was blinded to the genotype of the test mice throughout the experiment.

### Forced swim test

For the forced swim test, mice were habituated to the behavioral room 30 mins prior to the experiment. A 4L beaker was filled 2/3 of the way with water and was maintained at room temperature (24°C ± 1°C), measured before each experiment with a glass thermometer. Mice were gently placed in water and were closely observed throughout the 6-min test. Mice were immediately removed if they were unable to keep their head above water. After the test, mice were gently dried with a paper towel and were allowed to recover in the home cage for 10 mins under monitoring. Time immobile was manually scored with a stopwatch by an experimenter blinding to the animal genotype.

### Prepulse inhibition

For prepulse inhibition, mice were habituated to the behavioral room 30 mins prior to the start of the experiment. During the experiments, mice were allowed 5 mins to acclimate to the 60 db background noise that was present throughout the experiment. Mice were presented with 4 variable auditory pre-pulse levels (70, 74, 78, 82 dV; 20ms), followed by an auditory startle pulse (110 dB, 20ms) 100ms after the pre-pulse. A total of 20 trials is delivered for each mouse (5 repeats for each of the 4 pre-pulse levels) in a random order with a variable inter-trial-interval ranging from 10-30 seconds. The startle response was measured during the 250ms period after the onset of the startle pulse by a force meter (Kinder Scientific). The percentage of prepulse inhibition is quantified by the following equation: %PPI = 100 × [(pulse-alone) – (pre-pulse + pulse score)]/pulse-alone score.

### Three-chamber social interaction

Mice were habituated in the room 30 mins prior to the start of testing. A plastic apparatus (60 × 40 × 20 cm) with an open top divided into 3 chambers (20 cm × 40 cm × 20 cm) by 2 transparent walls was used. The testing session starts with a 10 min habituation period, during which chambers are separated by a wall containing a small circular opening. TEST mice are first placed in the middle chamber and are allowed to explore the entire arena. To examine social interaction after the habituation period, an unfamiliar, sex-matched mouse (STRANGER 1) is placed in one of the side chambers. The other side has an empty cage to control for object novelty.

STRANGER 1 is enclosed in a small round wire cage, which allows nose contact between the bars, but prevents fighting. The TEST mouse is allowed to explore the entire chamber during the 10 min session. The amount of time spent investigating the mice or the empty cage, time spent in each chamber, and the number of entries into each chamber are automatically scored using Ethovision. To examine social memory, after an interval of 10 min where the TEST mouse is returned to the home cage, the TEST mouse is tested in a second 10 min session to quantify social preference for a new stranger. In this second trial, STRANGER 1 remains in the wire cage and a second, unfamiliar mouse (STRANGER 2) is placed in an identical small wire cage located on the opposite side of the chamber. The TEST mouse now has a choice between the familiar mouse (STRANGER 1) and the novel mouse (STRANGER 2). The amount of time spent investigating each of the mice and the number of entries into each chamber is scored automatically using Ethovision.

### Accelerating Rotarod

A subset of male and female mice that underwent homecage assay (n= 7 WT, n=6 *Akap11^+/-^*, n=6 *Akap11^-/-^*) were evaluated with the rotarod test (Harvard 76-0770) when they were 6 months old. Biological sex was approximately evenly split between genotypes. Mice were first allowed to acclimate to the testing room in their home cages for at least 30 minutes prior to beginning testing. Mice underwent 3 training trials where they acclimated to walking on the rotating rod spinning at 4 rotations per minute (rpm). Two mice (n=1 *Akap11^+/-^*, n=1 *Akap11^-/-^*) were excluded from testing because they fell off immediately during all three training trials. Mice then underwent 4 test trials, with at least 1 minute between trials, where the rod accelerated at a constant rate from an initial speed of 4rpm until the mouse was unable to keep pace with the rotations and fell off of the rotating bar, or until a maximum speed of 40 rpm. This latency to fall was recorded and the data is shown as the average of 4 test trials.

### Anti-AKAP11 antibodies

Two custom polyclonal rabbit anti-AKAP11 antibodies were generated for this paper and used for immunoblot experiments. For both polyclonal antibodies, the first 284 amino acids of the mouse AKAP11 protein (Uniprot reference E9Q777) were injected into rabbits, and affinity purified by a company (Genscript). The antigen sequence used was: MAAFQPLRSSHLKSKASVRKSFSEDVFRSVKSLLQSEKELCSVSGGECLNQDEHPQLTEVTFLGFNEETDAAHIQDLAAVSLELPDLLNSLHFCSLSENEIICMKDTSKSSNVSSSPLNQSHHSGMLCVMRVSPTLPGLRIDFIFSLLSKYAAGIRHTLDMHAHPQHHLETTDEDDDDTNQSVSSIEDDFVTAFEQLEEEENAKLYNDEINIATLRSRCDAASQTTSGHHLESHDLKVLVSYGSPKSLAKPSPSVNVLGRKEAASVKTSVTTSVSEPWTQRSLY and were validated by immunoblot comparisons between WT and *Akap11^-/-^* brain tissue. For immunoprecipitations, we used a commercial rabbit monoclonal antibody targeting residues surrounding Alanine 1125 of mouse AKAP11 (Cell Signaling Technologies (CST), #77313). All other AKAP11 antibodies we tested, including our own polyclonal antibodies, immunoprecipitated PKA subunits in *Akap11^-/-^*brains, indicating insufficient specificity (data not shown but available upon request).

### Immunoprecipitation

Twelve week-old (P120) male mice were euthanized using CO_2_ inhalation, harvested around the lights-off period (Zeitgeber Time 12, ZT12) when mice are active, which was approximately ZT10-ZT14. For coIP-MS, 5 WT, 5, *Akap11^+/-^*, and 5 *Akap11^-/-^* animals underwent immunoprecipitation with the AKAP11 antibody. Three WT animals underwent immunoprecipitation with IgG control. After sacrifice, cortical hemispheres were rapidly dissected and mechanically homogenized with 12 slow strokes by a Teflon homogenizer and a glass vessel in ice-cold phosphate buffered saline (PBS) with 1% Triton-X with protease (Sigma 4693159001) and phosphatase (Sigma 4906837001) inhibitors. The homogenate was centrifuged at 10,000 g (4°C) for 10 minutes and the resultant supernatant was used for immunoprecipitation. Protein concentration was measured with bicinchoninic acid assay BCA (Thermo Fisher Scientific A53225) and samples were normalized to equal protein concentrations and volumes with homogenization buffer in protein LoBind tubes (Eppendorf #022431102). For coIP-mass spectrometry, samples had 4mg of protein in a volume of 2mL. For coIP-immunoblot, samples had either 500μg or 1mg (which were matched between experimental and control samples), normalized to 1 mg/mL. Lysates were incubated overnight rotating at 4°C with 1μg anti-AKAP11 monoclonal antibody (1:20; CST 77313) or 1μg mAb IgG control (CST 3900) for mass spectrometry experiments. For coIP-immunoblot, lysates were incubated with the following primary antibodies: 1:50 Rabbit anti-AKAP11 (CST 77313), 1:50 Rabbit anti-GluA1 (CST 13185), 1:50 Rabbit anti-PKA Cα (CST 4782), 1:50 Rabbit anti-PKA R1α (CST 5675), 1:50 Rabbit anti-PKA R2β (Sigma ABS14), 1:50 Rabbit anti-VAPA (Proteintech 15275-1-AP), 1:50 Rabbit anti-VAPB (Proteintech 14477-1-AP), 1:50 Rabbit anti-GSK3α (CST 4337), 1:50 Rabbit anti-GSK3β (CST 12456), 1:50 Mouse anti-DYRK1A (Abnova H00001859-M01), 1:50 Mouse anti-P62 (Abcam ab56416). As additional negative controls, Rabbit mAb IgG (CST 3900), Normal Rabbit IgG (CST 2729) or Mouse mAb IgG2a Isotype Control (CST 61656) were diluted according to manufacturer’s instructions and added in lieu of primary antibodies.

The next day, lysate-antibody mixtures were incubated with Dynabead Protein G (Invitrogen 1004D) for 1 hour at room temperature (RT), following the manufacturer’s specifications. Beads were then washed three times for 10 min with PBS containing 0.02% Tween20. For coIP-immunoblotting, proteins were eluted from magnetic beads with 1x Laemmli buffer and beta-mercaptoethanol (BME) mixture (Boston biosciences BP111R, Biorad 1610737, Sigma) and boiled at 98°C for 8 min and subsequently examined by SDS-PAGE (immunoblot). For coIP-MS, samples were washed twice with ice cold 50mM Tris-HCl pH 7.5 and transferred to a new tube. Subsequently samples were resuspended in 50μl ice cold Tris-HCl pH 7.5.

### CoIP sample preparation for Liquid Chromatography-Mass Spectrometry/Mass Spectrometry (LC-MS/MS) analysis

Samples were incubated in 80 μl of urea-based digestion buffer (30 mM Tris-HCl pH=8, 0.4 μg trypsin, 1mM dithiothreitol, 2 M Urea) at 25℃ and 1000 rpm for 1h. The supernatant was then collected, followed by a second incubation in 80 μl of the same urea-based buffer at 25℃ and 1000 rpm for 30 min. The second supernatant was also collected. The beads were washed with 60 μL of wash buffer (2M Urea, 50mM Tris-HCl pH=8 wash buffer) two times, collecting both washes. The on-bead digests (supernatants) and two washes were combined into 280 μl per sample and reduced with 2.3 μl of 500mM dithiothreitol (DTT) for 30 min at 1000 rpm at 25℃. This was followed by alkylation with 4.6 μl of 500 mM iodoacetamide (IAA) for 45 min at 1000 rpm at 25℃. The samples were digested overnight with 0.5 μg of Trypsin at 700 rpm at 25℃. Reactions were quenched with formic acid (FA) at a final concentration of 1%, then desalted using reverse-phase C-18 StageTips as previously described^50^ and dried down. The peptides were reconstituted in 80 μl 50mM HEPES, followed by 8 μl of neat acetonitrile (MeCN), and isobarically labeled using 500 μg of Tandem Mass Tag 18-plex (TMT18) reagent (Thermo Fisher Scientific A44520 & A52046) for 1h at 1000 rpm and 25℃. After confirming successful labeling ( > 95% label incorporation), the reactions were quenched with 4 μl of 5% hydroxylamine. Samples were mixed and dried. The sample was resuspended in 300 μl of 0.1% formic acid to proceed with fractionation by bRP fractionation strategy on StageTips. The tips were packed with 3 plugs of Empore Extraction Disks SDB-RPS 47mm material. StageTips were sequentially conditioned with 100 μl of methanol (MeOH) and 100 μl of 50%MeCN/0.1%FA. The sample was loaded onto the StageTips, and washed twice with 100 μl of 0.1% FA. 25 μl of 20 mM ammonium formate was added to the stage tip for buffer exchange. It was then eluted into 8 consecutive fractions with increased organic content at 100 μl per elution step (5, 10, 15, 20, 23, 27, 30 and 45% MeCN in 20 mM ammonium formate). The fractions were transferred to HPLC vials and dried.

### LC-MS/MS analysis of coIP samples

The samples were analyzed on a Q Exactive HF-X mass spectrometer with a Vanquish Neo UHPLC system (Thermo Fisher Scientific). The solvents used for the HPLC system were solvent A (0.1% FA) and solvent B (0.1% FA/100% MeCN). The samples were injected into a 75-μm ID PicoFrit column packed manually to approximately 28 cm of length with ReproSil-Pur C18-AQ 1.9-μn beads (Dr. Maisch) and heated to 50℃ for chromatographic separation.

The bRP fractions were loaded into the column at a flow rate of 200nL/min. The chromatography consists of a 110-min method with a gradient of 1.6–5.4% solvent B for 1 min, 18–27% B for 22 min, 27–54% B for 9 min, 54–72% B for 1 min, followed by a hold at 72% B for 5 min and 72%-45% B for 10 min. The mass spectrometer was operated in data-dependent acquisition mode. The MS parameters were set as follows: MS1, r=60,000; MS2, r=15,000; MS1 AGC target of 3e6 with maximum injection time of 10ms; MS2 for the 20 most abundant ions using an AGC target of 5e4 and maximum injection time of 105ms; and normalized collision energy of 28.

### Data analysis of the coIP-LC-MS/MS experiment

Collected data were analyzed using the Spectrum Mill software package (Broad Institute). All extracted spectra were searched against a UniProt database containing mouse reference proteome sequences retrieved on 04/07/2021 and containing 55734 entrees. Fixed modifications were carbamidomethylation at cysteine. TMT labeling was required at lysine, but peptide N termini could be labeled or unlabeled. Allowed variable modifications were protein N-terminal acetylation and oxidized methionine.

Protein quantification was achieved by taking the ratio of TMT reporter ions for each sample over the median of all channels. TMT18 reporter ion intensities were corrected for isotopic impurities in the Spectrum Mill protein/peptide summary module using the afRICA correction method which implements determinant calculations according to Cramer’s Rule79 and correction factors obtained from the reagent manufacturer’s certificate of analysis for lot numbers XD343709 and XD346675. After performing median-MAD normalization, a moderated two-sample t-test was applied to the datasets to compare WT, *Akap11^+/-^* and *Akap11^-/-^* sample groups.

### Immunoblot

Four week (P30) or 12 week (P120) male mice were euthanized using CO_2_ inhalation between ZT10-ZT14, after which whole cortices were rapidly dissected and flash-frozen in liquid nitrogen. Single hemispheres were homogenized in a 5 ml ice-cold homogenization buffer (1% SDS in PBS) supplemented with phosphatase (Sigma 4906837001) and protease inhibitors (Sigma 4693159001), using the mechanical homogenization procedure described above. After BCA (Thermo Fisher Scientific, A53225 or 23225) quantification of protein concentration, samples were diluted with 6X Laemmli buffer with BME (Boston Bioproducts, BP-111R) or 2x Laemmli buffer (Biorad, 1610737) with BME (Sigma, M3148) to a final concentration of 1x and boiled in reducing SDS loading buffer at 98°C for 8 min. After preparation, sample aliquots were frozen at -80°C. Before running on a gel, samples were thawed, heated at 98°C for 1 min and centrifuged briefly. Samples were run on NuPAGE 3-8% Tris Acetate gradient gels (Invitrogen) at a constant voltage. Equal amounts of sample were loaded unless otherwise specified (Supplementary Fig. 4a,b). For Fig. 3a, a subset of smaller molecular weight proteins were tested on 4-20% tris glycine gradient gels (Invitrogen). Proteins were transferred to 0.2 mm nitrocellulose membranes using semi-dry transfer (Bio-Rad Transblot Turbo; 25V 30 min) and visualized by Coomassie stain (Sigma, P7170). Subsequently, membranes were blocked with 5% milk in Tris-buffered saline with 0.2% Tween 20 (TBST). Membranes were incubated overnight with primary antibodies in 1% milk TBST with gentle rotation at 4°C. Membranes were incubated with primary antibodies overnight at 4°C, washed three times with TBST for 10 min, and then incubated with secondary HRP-conjugated antibodies raised against the appropriate species for 1 hour at RT. After 3 additional TBST washes, membranes were imaged on ChemiDoc MP (Bio-Rad) and analyzed using Image Lab Software (Bio-Rad). Enhanced chemiluminescence (Thermo Fisher Scientific, 34580, 34096 and A38556) and exposures were optimized to obtain signals in the linear range. Membranes were then stripped (Thermo, 46430),washed three times, re-blocked with 5% milk in TBST, and re-probed as described above. For cortical lysate immunoblots, 20μg of protein was loaded. For microdissected brain regions, 15 or 20 μg of protein and 5 or 10 μg synapse fraction protein were used for SDS-PAGE, unless otherwise specified (Supplementary Fig. 4a,b). For all experiments, protein loading amounts were consistent across samples.

For coIP-immunoblot, the following antibodies were used during SDS-page (after immunoprecipitation): 1:1000 Mouse anti-PKA R1α (BD 610609), 1:1000 Mouse anti-PKA Cα (BD 610980), 1:1000 Mouse anti-PKA R2β (BD 610625), 1:1000 Mouse anti-GluA1 (NeuroMab 75-327), 1:1000 Rabbit anti-Vinculin (CST 4650), 1:1000 Rabbit anti-VAPA (Proteintech 15275-1-AP), 1:1000 Rabbit anti-VAPB (Proteintech 14477-1-AP), 1:1000 Rabbit anti-GSK3α (CST 4337), 1:1000 Rabbit anti-GSK3β (CST 9315), Mouse anti-GSK3α/β (Thermo Fisher Scientific 44-610) ,1:1000 Rabbit anti-Dyrk1a (CST 2771), 1:1000 Rabbit anti-P62 (CST 5114). For coIP-immunoblot, we used horseradish peroxidase (HRP)-conjugated secondaries recognizing the host species of the primary antibody (Thermo Fisher Scientific 31460, 31430 and A18775 for rabbit, mouse and guinea pig respectively). When the IP and immunoblot antibodies were from the same species (Fig. 2g), we used secondaries that are selective towards non-denatured IgG (1:1000, TrueBlot HRP, Rockland, 18-8816-33 and 18-8817-30).

For immunoblot, the following antibodies were used: 1:1000 Rabbit anti-AKAP11 (Custom R171), 1:1000 Rabbit anti-AKAP11 (Custom R173), 1:1000 Mouse anti-PKA R1α (Thermo Fisher Scientific MA5-24981), 1:1000 Rabbit anti-PKA R1α (CST 5675), 1:1000 Rabbit anti-PKA Cα (CST 4782), 1:1000 Rabbit anti-Iqgap1 (Abcam ab86064), 1:1000 Rabbit anti-Dyrk1a (CST 2771), 1:1000 Rabbit anti-GSK3α (CST 4818), 1:1000 Rabbit anti-GSK3β (CST 9315), 1:1000 Mouse anti-P62 (Abcam ab56416), 1:1000 Rabbit anti-LC3A/B (CST 12741), 1:1000 Rabbit anti-VAPA (Proteintech 15275-1-AP), 1:1000 Rabbit anti-β Catenin (CST 8480), 1:1000 Mouse anti-Protein Phosphatase 1 (Santa Cruz SC-7482), 1:1000 Rabbit anti-Sik2 (CST 6919), 1:1000 Rabbit anti-p-GSK3α^S21^ / anti-p-GSK3β^S9^ (CST 8566), 1:1000 Rabbit anti-p-GSK3α / anti-p-GSK3β (CST 5676), 1:1000 Rabbit anti- p- GSK3β^S9^ (CST 9336), 1:1000 Rabbit anti- p-GluA1^S845^ (CST 8084), 1:1000 Rabbit anti- GluA1 (CST 13185), 1:1000 Mouse anti- GluA1 (NeuroMab 75-327), 1:1000 Mouse anti-PSD95 (BioLegend 810301), 1:1000 Rabbit anti-Homer1 (Synaptic Systems 160003), 1:1000 Guinea Pig anti-Synaptophysin 1(Synaptic Systems 101004). 1:1000 Mouse anti-PSD95 (clone K28/43, Antibodies Inc 75-028). HRP-conjugated anti-Rabbit or anti-Mouse IgG (Thermo Fisher Scientific 31460 and 31430). β-actin was probed using an HRP conjugated primary antibody after all other proteins were probed (1:10,000. Thermo Fisher Scientific A3854). Otherwise, we used HRP conjugated secondaries recognizing the host species of the primary antibody (Thermo Fisher Scientific).

### Synapse purifications

Synapse fractions were purified as we described previously^50,116,117^. Flash-frozen whole cortex with subcortical forebrain was mechanically homogenized with 12 slow strokes by a Teflon homogenizer and glass vessel in ice-cold homogenization buffer (5 mM HEPES pH 7.4, 1 mM MgCl_2_, 0.5 mM CaCl_2_, supplemented with phosphatase (Sigma 4906837001) and protease inhibitors (Sigma 4693159001). The homogenate was centrifuged for 10 min at 1,400 g (4°C) and the resulting supernatant (S1) was re-centrifuged at 13,800 g for 10 min (4°C). The resulting pellet (P2) was resuspended in 0.32 M Sucrose, 6 mM Tris-HCl (pH 7.5) and layered gently on a 0.85 M, 1 M, 1.2 M discontinuous sucrose gradient (all layers in 6 mM Tris-HCl pH 7.5) and ultracentrifuged at 82,500 g for 2 hours (4°C). For synapse mass spectrometry experiments, phosphatase inhibitors were added to 1 M and 1.2 M sucrose buffers. The synaptosome fraction, which sediments at the interface between 1 M and 1.2 M sucrose, was collected. An equal volume of ice-cold 1% Triton X-100 (in 6 mM Tris-HCl pH 7.5) was added, mixed thoroughly, and incubated on ice for 15 min. The mixture was ultracentrifuged at 32,800 g for 20 min (4°C), and the pellet (the synapse fraction) was collected by resuspension in 1% SDS. A small aliquot was taken to measure the protein concentration using the Pierce Micro BCA assay (ThermoFisher Scientific, 23235) or BCA (ThermoFisher Scientific A53225) and the remaining protein was stored at -80°C until processed for MS or immunoblot. For synapse proteomics, sixteen synaptic fractions extracted from the cortex of wild type, *Akap11^+/-^* and *Akap11^-/-^* mouse brains were used for the proteomics study using tandem mass tag (TMT) isobaric labeling strategy for quantitation. At 4 weeks, sample sizes were n= 5 WT, 5 *Akap11^+/-^* and 6 *Akap11^-/-^*. At 12 weeks, the sample sizes were n= 6 WT, 5 *Akap11^+/-^* and 5 *Akap11^-/-^*.

### Processing of samples for proteomics and phosphoproteomics

Synapse fractions from animals at 4 weeks and 12 weeks were processed in two separate TMT16 plexes as described before^50^. Briefly, all samples were prepared in a 1% SDS buffer and digested using S-Trap^TM^ sample processing technology (Protifi) following manufacturers recommendations. Following digestion, 100 μg of each sample was reconstituted in 20μl of 50mM HEPES and labeled with 200 μg TMT16 reagent (Thermo Fisher Scientific A44520).

After verifying successful labeling of more than 95% label incorporation reactions were quenched, mixed, desalted and dried down. TMT16 labeled peptides were fractionated by high pH reversed-phase chromatography on a 4.6 mm x 250 mm Zorbax 300 extend-c18 column (Agilent). One-min fractions were collected during the entire elution and fractions were concatenated into 18 proteome fractions for LC-MS/MS analysis. Five percent of each fraction was removed for proteome analysis. The remaining of each of the 18 fractions were further concatenated to 9 fractions and dried down for phosphopeptide enrichment using immobilized metal affinity chromatography (IMAC).

### LC-MS/MS analysis for synapse fractions

One microgram of each proteome fraction and half of each of the IMAC fractions were analyzed on a QE-HFX mass spectrometer (Thermo Fisher Scientific) coupled to an Easy-nLC 1200 LC system (Thermo Fisher Scientific). Samples were separated using 0.1% Formic acid/3% Acetonitrile as buffer A and 0.1% Formic acid /90% Acetonitrile as buffer B on a 27cm 75um ID picofrit column packed in-house with Reprosil C18-AQ 1.9 mm beads (Dr Maisch GmbH) with a 110 min gradient consisting of 2-6% B in 1 min, 6-20% B in 62 min, 20-30% B for 22 min, 30-60% B in 9 min, 60-90% B for 1 min followed by a hold at 90% B for 5 min. The MS method consisted of a full MS scan at 60,000 resolution and an AGC target of 3e6 from 350-1800 m/z followed by MS2 scans collected at 45,000 resolution with an AGC target of 3e6 from 300-1800 m/z followed by MS2 scans collected at 35,000 resolution with an AGC target of 1e5 and 5e4 for proteome and phosphoprotome fractions, respectively and a maximum injection time of 105 ms. Dynamic exclusion was set to 15 seconds. The isolation window used for MS2 acquisition was 0.7 m/z and 20 most abundant precursor ions were fragmented with a normalized collision energy (NCE) of 29 optimized for TMT16 data collection.

### Database search and MS/MS quantification for PSD fractions

All the datasets were searched on Spectrum Mill MS Proteomics Software (Broad Institute) using mouse database downloaded from Uniprot.org on either 12/28/2017 or 04/07/2021 containing 47,069 or 55,734 entrees for 4 weeks and 12 weeks coIP-MS experiments, respectively^118^. Search parameters included: ESI Qexactive HCD v4 35 scoring parent and fragment mass tolerance of 20 ppm, 40% minimum matched peak intensity, trypsin allow P enzyme specificity with up to four missed cleavages, and calculate reversed database scores enabled. Fixed modifications were carbamidomethylation at cysteine. TMT labeling was required at lysine, but peptide N-termini could be labeled or unlabeled. Variable modifications were allowed for protein N-terminal acetylation and oxidized methionine as well as phosphorylated serine, threonine, and tyrosine for the phosphoproteome datasets. Protein quantification was achieved by taking the ratio of TMT reporter ions for each sample over the TMT reporter ion for the median of all channels. TMT16 reporter ion intensities were corrected for isotopic impurities in the Spectrum Mill protein/peptide summary module using the afRICA correction method which implements determinant calculations according to Cramer’s Rule and correction factors obtained from the reagent manufacturer’s certificate of analysis (Thermo Fisher Scientific, 90406) for lot numbers VH310017 for both synapse proteomics experiments. Median normalization was applied to both proteome and phosphoproteome datasets prior to statistical analysis. Additionally, phosphosite changes were normalized for changes in protein abundances by fitting a global linear model between the phosphosite and the cognate protein and extracting the residuals. A moderated two-sample t-test was applied to the datasets to compare WT, *Akap11^+/-^* and *Akap11^-/-^* sample groups. For proteins with multiple isoforms, the isoform with the greatest abundance (number of spectra observed) was displayed in the plot.

### Annotations from Phosphosite Plus

Kinase substrate annotations were downloaded from https://www.phosphosite.org on June 20, 2022. In this database, some kinase substrate sites are inaccurately attributed to a kinase to which it is indirectly related. For example, Grin2b^Y1472^ is annotated as a PKA Cα substrate in phopshopsite.org (see Fig. 5b, but is actually a PKC substrate^119^. Similarly, Ctnnb1^S45^ is annotated as a Gsk3β substrate in phosphosite.org, but is actually a CK1 substrate^57^. These discrepancies were not adjusted for this manuscript in order to maintain unbiased usage of the database.

### Phosphoproteomics motif analysis

Motif enrichment analysis was performed to generate motif logos using the pyproteome Python library^120^. Motif logo enrichment values were calculated according to the method described for pLogo^121^. Peptide 15-mers around each phosphorylation site were generated by Spectrum Mill “VMsiteFlanks”. Foreground and background N-mers were extracted from protein-aligned sequences, using ‘-’ to pad N-and C-terminal peptides. Quantification values from identical N-mers (mis-cleaved, oxidized, or multiply-phosphorylated peptides) were combined by taking the median value. Log odds enrichment values are equal to 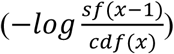, with k = the count of amino acids at that relative position for phosphosites in a foreground collection (*P* < 0.05; FC > 1.25 or FC < 0.8). Survival function (sf) and cumulative distribution functions (cdf) were calculated using scipy’s hypergeometric distribution model (scipy.stats.hypergeom), with: *n* = the total number of foreground N-mers; K = the count of amino acids at that position in the background collection; and *N*=the total number of background N-mers.

### PKA activity assay

12 week) male mice were euthanized using CO_2_ inhalation between ZT10-ZT14. Cortex and subcortical forebrain were separated during dissection, and flash-frozen in liquid nitrogen. The subcortical forebrain is enriched for the striatum but also includes other regions such as basal forebrain and hypothalamus. For this experiment only, we refer to the subcortical forebrain as “Striatum”. This crude dissection was necessary since the synapse-enrichment protocol requires large amounts of starting material. Single cortical hemispheres and both combined subcortical forebrains from both hemispheres were prepared as described in the “synapse purifications” section. Total lysate and synapse fractions were collected from the same starting material and resuspended in 1% Triton-X-100 and frozen on dry ice and stored at -80. After all samples were collected, samples were thawed and protein concentration was measured using Pierce Micro BCA assay. A colorimetric PKA activity assay (Invitrogen EIAPKA) was conducted according to the manufacturers instructions. 50ng of cortex and striatum samples were compared in separate plates. Lysate and synapse fractions were compared within the same plate. Curve-fitting functions in Graphpad Prism were used to infer PKA activity from provided standards.

### Quantitative Immunofluorescence

15-17-week-old male mice were euthanized under isoflurane with transcardial perfusion of ∼25 mL with PBS followed by ∼25 ml 4% paraformaldehyde (PFA). These mice also underwent an 1 hour open field assays for a separate study^14^, at 4 weeks and 12 weeks of age. Brains were post-fixed overnight in 4% PFA, and washed the next day four times with PBS for 15 min. Tissues were cryoprotected for 72 hours and stored at 4°C until sectioning. Brains were sectioned on a vibratome (Leica) at either 40 or 60μm thickness (consistent within technical replicates) and stored in a freezing buffer (0.1M PBS, 30% ethylene glycol, 30% glycerol, 0.01% sodium azide).

For immunostaining, free-floating sections were first rinsed 4x with PBS for 10 min. Sections were blocked and permeabilized with 1 hour RT incubation in a blocking solution. For the first technical replicate, we used 5% Normal Goat Serum (Sigma, NS02L), with 10% BSA + 0.2% Triton. For the second technical replicate, we used 5% Normal Donkey Serum (Jackson, 017-000-121) + 0.2% Triton in PBS for 60 min. Primary antibodies were diluted in blocking solution and incubated overnight at 4°C, rotating gently. The following day, sections were rinsed 3x for 15 min in PBS containing 0.2% Triton at RT on a shaker. DAPI (Thermo Fisher Scientific, 62248) was added to PBS at a final concentration of 1:5000 (for the first technical replicate) or 1:10,000 (for the second technical replicate) before mounting. The following primary and secondary antibodies were used: PKA Ca (CST 4782, 1:100), PKA R1a (ThermoFisher MA5-24981, 1:100). Donkey anti Rabbit 488 (Invitrogen A32790), Donkey anti-Mouse 555 (Invitrogen A32773), Goat anti Rabbit 488 (Invitrogen A11034), Goat anti-Mouse 555 (Invitrogen A32727). Whole sections were imaged with a Zeiss Axioscan 7 Slidescanner using exposure and intensity settings that were constant across samples within technical replicates.

PKA R1α was quantified in FIJI using the freehand selection tool and the intensity measurement function. Regions were selected using the Allen Brain Atlas as a reference. Experimenters were blind to conditions during analysis. For representative images, brightness levels were uniformly increased across the whole images with Zen software (Carl Zeiss) and scaled down to 10% to accommodate the large field of view of the stitched image.

### Tissue collection for RNA sequencing

For bulk and snRNA-seq, mice were harvested between ZT9 and ZT15, near the animals active period at ZT12. Brain tissues for bulk and snRNA-seq were prepared as described^12^, and also available at protocols.io (https://www.protocols.io/view/fresh-frozenmouse-brain-preparation-for-single-nu-bcbrism6). At 4 and 12 weeks of age, male WT, *Akap11^+/-^* and *Akap11^-/-^* mice were anesthetized by administration of isoflurane in a gas chamber. Within each age, 4 mice per genotype were harvested for snRNA-seq, with the exception of 4 week striatum, which had 5 WT samples. For bulk RNA-seq, 5-6 mice per genotype within an age were collected, with the exception of 4 week *Akap11^+/-^*hippocampus, which had 4 samples.

Transcardial perfusions were performed with ice-cold Hank’s Balanced Salt Solution (HBSS, Gibco 14175-095) in mice anesthetized with 3% isoflurane delivered continuously via nose cone. Brains were then frozen gradually over liquid nitrogen vapor and stored at –80°C with tubes containing O.C.T. (optimal cutting temperature) freezing medium (Tissue-Tek) to prevent desiccation. Brain region microdissections were performed by hand in a cryostat (Leica CM3050S), with a ophthalmic microscalpel (Feather safety razor P-715) or 1.5 mm diameter biopsy punch (Integra 33-31A). PFC, Hipp and SSC were excised by hand. SN and Thal were collected by biopsy punch. For snRNA-seq studies, both dorsal and ventral striatum were collected together with a microscalpel, while for bulk RNA-seq, dorsal striatum was collected by punch. All brain regions were identified according to the Allen Brain Atlas. Dissected tissues were placed into a precooled 1.5-ml PCR tube and stored at –80°C.

### RNA extraction and bulk RNA-seq library preparation

RNA was prepared from micro-dissected tissues using RNeasy Mini Kit (Qiagen, 74106) following the manufacturer’s instructions. Briefly, tissue samples were lysed and homogenized, placed into columns, and bound to the RNeasy silica membrane. Next, contaminants were washed away. Columns were treated with DNase (Qiagen) to digest residual DNA and concentrated RNA was eluted in water. RNA concentration was measured using a NanoDrop Spectrophotometer and RNA integrity (RIN) with RNA pico chips (Agilent) using a 2100 Bioanalyzer Instrument (Agilent). Purified RNA was stored at -80°C until library preparation for bulk RNA-seq analysis. Bulk RNA-seq libraries were prepared using a TruSeq Stranded mRNA Kit (Illumina) following the manufacturer’s instructions. 200 ng of isolated total RNA from each sample was used and the concentration of resulting cDNA library was measured with High Sensitivity DNA chips (Agilent) using a 2100 Bioanalyzer Instrument (Agilent). A 10 nM normalized library was pooled, and sequencing was performed on a NovaSeq S2 (Illumina) with 50 bases each for reads 1 and 2 and 8 bases each for index reads 1 and 2.

For snRNA-seq library preparation, brain tissue dissection and nuclei extraction were performed within 16 hours to avoid desiccation and freeze-thaw cycles. A gentle, detergent-based dissociation was used to extract the nuclei, according to a previously published protocol^122^ also available at protocols.io (https://www.protocols.io/view/frozen-tissue-nuclei-extraction-bbseinbe). Extracted nuclei were then loaded into the 10x Chromium V3.1 system (10x Genomics) and library preparation was performed according to the manufacturer’s protocol. A 10 nM normalized library was pooled, and sequencing was performed on a NovaSeq S2 (Illumina) with 28 and 75 bases for reads 1 and 2, respectively, and 10 bases each for index reads 1 and 2.

### Bulk RNA-seq DE analysis

FASTQ files were aligned to a reference, which was derived from the genome FASTA file and transcriptome GTF extracted from the CellRanger mm10 reference. Alignment and quantification were performed using a Nextflow V1 pipeline^123^ (https://github.com/seanken/BulkIsoform/blob/main/Pipeline). Quantification was performed by Salmon^124^ (version 1.7.0, with arguments -l A --posBias --seqBias --gcBias --validateMapping), using a Salmon reference built with the salmon index command with genomic decoys included. Two samples - HP_HT_941 and TH_WT_913 - were removed from analysis due to evidence of contamination by aberrant marker gene expression.

| Dataset | WT sample size (*n*) | *Akap11*<sup>+/-</sup> sample size (*n*) | *Akap11*<sup>-/-</sup> sample size (*n*) |
| --- | --- | --- | --- |
| 4wk PFC (Prefrontal Cortex) | 5 | 6 | 5 |
| 4wk Hipp (Hippocampus) | 6 | 4 | 5 |
| 4wk SSC (Somatosensory Cortex) | 5 | 6 | 5 |
| 4wk Striatum (Str) | 5 | 5 | 5 |
| 4wk Thalamus (Thal) | 5 | 5 | 5 |
| 4wk Substantia Nigra (SN) | 6 | 6 | 5 |
| 12wk PFC | 5 | 5 | 5 |
| 12wk Hipp | 5 | 5 | 5 |
| 12wk SSC | 5 | 5 | 5 |
| 12wk Str | 5 | 5 | 5 |
| 12wk Thal | 5 | 5 | 5 |
| 12wk SN | 5 | 5 | 5 |

### Differential expression

Differential expression analysis was performed to compare the heterozygous or knockout mutants to the wild-type mice for each brain region and age using the Wald test in R package DESeq2^125^ (version 1.34). Salmon output was loaded using R package tximport^126^ (version 1.22). Genes that had less than 10 counts across all samples in each experiment were removed from our analysis. Due to evidence of dissection variability in striatum by excitatory marker genes, DESeq2’s PC1 was added as a covariate for differential expression in 1 month striatum. Log2 fold change shrinkage was applied using the “normal” shrinkage estimator in DESeq2. DESeq2’s default results filtering was used and all genes in a given comparison with padj = NA were removed from downstream analysis. For 1 month hippocampus and thalamus, the default filtering removed the majority of genes, so we changed the filtering for the heterozygous vs. WT comparison to match the less aggressive filtering in the knockout vs. WT comparison. The statistical method applied for bulk RNAseq is two-tailed Wald test, with Benjamini-Hochberg (B-H) false discovery rate (FDR) correction.

### Single-nucleus RNA-seq analysis

snRNA-seq reads were aligned to an mm10 mouse reference genome containing introns (https://cf.10xgenomics.com/supp/cell-exp/refdata-gex-mm10-2020-A.tar.gz) using the Cell Ranger v3.0.2 pipeline (10x Genomics)^127^. Cell Ranger count was run with default parameters and the following additional flags: --chemistry=SC3Pv3 and --expect-cells=15000. Replicates for the 4-week STR dataset were down-sampled to contain the same number of reads per cell using Cell Ranger aggr with default parameters.

Seurat v4.0.3^128^ was used to analyze Unique Molecular Identifier (UMI) counts. Nuclei expressing less than 500 genes were removed and the remaining nuclei were normalized by total expression, multiplied by ten thousand, and log transformed. The data were then scaled using Seurat’s ScaleData function. Seurat’s RunPCA function was used to perform linear dimension reduction on the variable genes within the scaled data. Clustering in the PCA space was then run using Seurat’s FindNeighbors and FindClusters functions and the clusters were visualized using Seurat’s RunUMAP function. Doublet identification was performed using the Scrublet v0.2.3^129^. Python package with default parameters. Clusters and nuclei with a scrublet score above the region-specific thresholds were determined to be doublets and were removed from the data. The table below shows the used scrublet score thresholds for all snRNA-seq datasets:

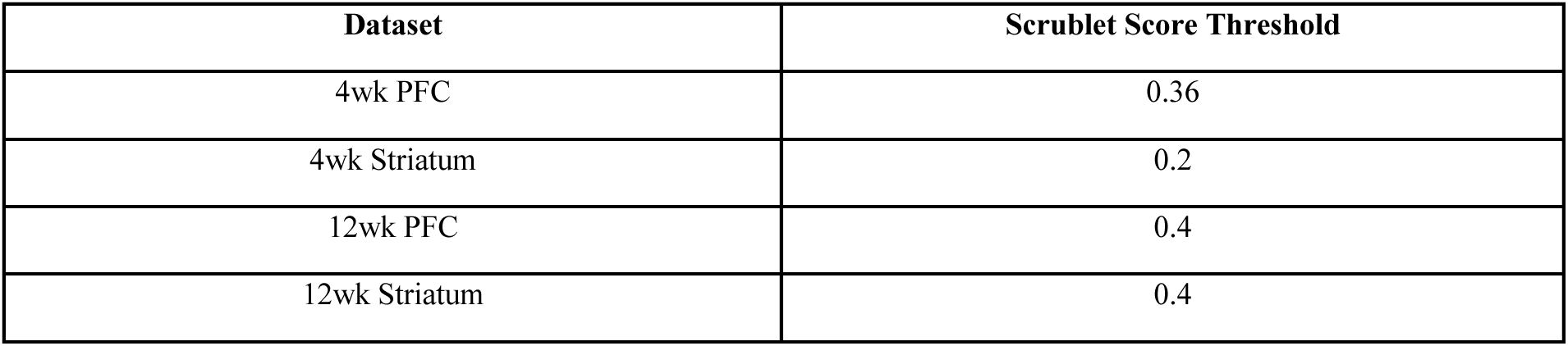

| Dataset | Scrublet Score Threshold |
| --- | --- |
| 4wk PFC | 0.36 |
| 4wk Striatum | 0.2 |
| 12wk PFC | 0.4 |
| 12wk Striatum | 0.4 |

Cluster cell type assignment was evaluated using region-specific cell type marker genes. Neuronal subtypes were identified by re-clustering the nuclei of a given major cell type and evaluating expression of subtype-specific marker genes. Marker genes for the more widely classified major cell types and subtypes were identified in the literature^130^. Rarer cell types and subtypes were identified using Seurat’s FindMarkers function. The clusters were then identified by searching cluster-specific marker genes on mouseBrain.org^131^ and or dropviz.org^132^.

Sample size for single-nucleus RNA-seq datasets:

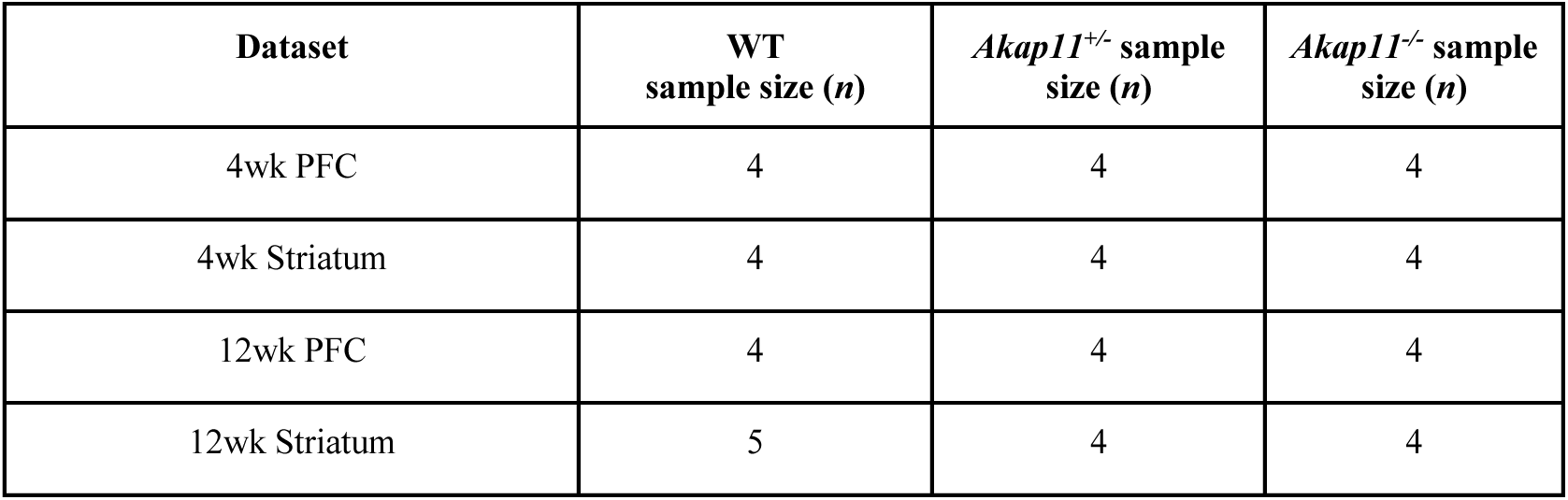

### Pseudobulk DE analysis

snRNA-seq differential expression (DE) analysis was performed using the pseudobulk approach. For each dataset, counts were grouped by cell type and replicate, then gene counts were summed across each replicate within a cell type, resulting in one count value per replicate per cell type.

Surrogate variable analysis (SVA) was conducted using the sva v3.42.0 R package^133^ to account for unknown sources of variation within each experiment. Differential expression analysis was then run with the covariates identified by SVA using the EdgeR v3.22.3 R package^134^.

Comparisons were calculated using two-tailed likelihood ratio test, which were FDR corrected with B-H adjustment. DEGs were defined as genes with a B-H adjusted *P*-value <0.05.

### Gene set enrichment analysis

GSEA was performed with the R Bioconductor package *fgsea*^135^ using C5 ontology gene sets obtained from the Molecular signatures database (MSigDB) on September 28, 2022. Genes were pre-ranked using their test-statistics within each comparison, and their *Homo sapiens* homolog-associated were used for the analysis. For genes or proteins with multiple isoforms, the isoform with the largest effect-size (absolute value of the t-statistic) was used for pre-ranking. Because the enrichment analyses were carried out using the proteins’ gene symbols, we use the terms gene set and protein set interchangeably in this study. For each of these gene sets, mouse gene symbols from transcriptomics and proteomics results were mapped to matching *Homo sapiens* homolog-associated gene symbols through annotations extracted from Ensembl’s BioMart data service; the human symbols were then used to perform GSEA. GSEA uses a Kolmogorov-Smirnov test, two tailed, with B-H FDR correction applied. A list of all significant GO terms (FDR < 0.05) for all datasets are provided as Supplementary Data. GSEA analysis with custom gene sets was conducted by appending these sets to the C5 MSigDB database.

### Curated gene sets

Phosphosites that correlate with homeostatic synaptic scaling were curated from Desch et al.^72^, including together all phosphosites that were significantly altered (adjusted *P*-value < 0.05) after 5 minutes, 15 minutes or 24 hours of either bicuculline or tetrodotoxin treatment. Curated gene sets that we have previously described^12^ include “Activity-regulated genes” (ARGs), which include rapid, delayed and slow primary response genes induced by neuronal depolarization *in vitro*^79^, GWAS and exome associations for neuropsychiatric traits^37,82–84^, and dopamine-induced genes which are genes with FDR < 0.1 that were upregulated in striatal neuronal culture following 1 hour dopamine treatment^136^.

### Brain slice electrophysiology

6-8-week-old male and female mice were used for brain slice electrophysiology experiments, including n = 9 slices from 4 mice WT, and n = 8 slices from 4 mice *Akap11^-/-^* for input-output experiments; n = 25 slices from 8 mice WT, n = 24 slices from 7 mice *Akap11^+/-^*, and n = 23 slices from 8 mice *Akap11^-/-^* for early LTP and paired-pulse facilitation experiments. An NMDG-based cutting solution was used and contains (in Mm): 93 NMDG, 2.5 KCl, 1.2 NaH_2_PO_4_, 30 NaHCO_3_, 20 HEPES, 25 Glucose, 10 MgSO_4_, 0.5 CaCl_2_, 5 Sodium Ascorbate, 2 Thiourea, 3 Sodium Pyruvate. pH = 7.3-7.4, OSM = 300-310. The external artificial cerebrospinal fluid (ACSF) recording solution contains(in Mm): 124 NaCl, 2.5 KCl, 1.25 NaH_2_PO_4_, 24 NaHCO_3_, 5 HEPES, 12.5 Glucose,1 MgSO_4_, 2 CaCl_2_ pH = 7.35, OSM = 300. For brain slice preparation, cardiac perfusion was performed with cold NMDG cutting solution under isoflurane anesthesia. Transverse (horizontal) brain slices containing hippocampus were cut at 400μm with a Leica VT1200S Microtome when submerged in ice-cold NMDG cutting solution. Brain slices were transferred to NMDG solution at 32°C for approximately 10 min before being transferred to RT brain slice holding solution. Slices were allowed to recover for at least 1 hour before starting the recordings.

To obtain hippocampus field electric postsynaptic potential (EPSP), a concentric bipolar stim electrode (FHC Inc) and a borosilicate glass recording electrode (2-3mOhm resistance) were placed in the classic schaffer collateral to CA1 (SC-CA1) pathway. The signals were low-pass filtered at 2.8kHz using a MultiClamp 700B amplifier (Molecular Devices) and digitized at 10kHz using a DAQ board (National Instrument). Baseline field EPSP were evoked by a paired-pulse stimulation (40ms ISI) every twenty seconds, and a stimulation intensity that evoked 40% of the maximum field EPSP response was used for baseline recording.

For input-output recordings, field EPSP responses were evoked with varying stimulation intensities (20-500 μA). For early LTP recordings, slices were stimulated for approximately 30 mins until a stable baseline field EPSP was obtained. Theta burst stimulation consists of short bursts of 4 stimulation pulses at 10Hz delivered at theta frequency (5Hz, 10 epochs) were used to induce LTP. The field EPSP was recorded for another 20 mins after LTP induction. Paired-pulse ratio was quantified using the field EPSP collected from the 10-minute period before LTP induction and was defined as the ratio between the second and the first field EPSP slope. Field EPSP slope was quantified as the slope during 10-90% of the raise time.

### Single-molecule fluorescent in-situ hybridization (smFISH)

Mouse brain characterization of *Akap11*, *Prkaca*, and *Prkar1a* expression was performed using Advanced Cell System’s (ACD) single-molecule *in situ* hybridization multiplexed fluorescent amplification kits (ACD catalog numbers 323270). 12-week-old male mice, n=6 per genotype WT, *Akap11^+/-^,* and *Akap11^-/-^*, were deeply anesthetized under isoflurane, perfused with ice-cold HBSS for 3 minutes, and decapitated. Brains were extracted and frozen gradually over liquid nitrogen vapors. Frozen brains were stored in tubes with O.C.T. in -80°C. Brains were sectioned sagittally at 12um/section using a Leica cryostat. Four sections per animal (two sections per slide) were collected on SuperFrost Charged slides and stored at -80°C and smFISH was performed the following day. Slides were post-fixed in freshly made, pre-chilled 4% PFA (Electron Microscopy Sciences) in PBS for 1 hour on ice. Sections were then dehydrated with 50%, 70%, and 100% ethanol for 5 min at room temperature. All slides were outlined with a grease pen and all following smFISH steps were done on a humidified slide rack. Sections were then treated with hydrogen peroxide for 10 min followed by three PBS washes. Sections then underwent permeabilization with “protease IV” for 30 min followed by three PBS washes.

Sections were then hybridized using probes to *Prkaca* (channel 1), *Akap11* (channel 2), and *Prkar1a* (channel 3) at 40°C for 2 hours, washed twice with wash buffer, and placed in aluminum foil-wrapped slide holders filled with 5X SSC (Thermo Fisher Scientific, 15557044) overnight at room temperature. The following day, sections were thoroughly washed in wash buffer and probes were amplified using the AMP reagents from the ACD kit. Slides were washed between each step using the wash buffer. HRP binding was also carried out according to the kit’s protocol with C1, C2, and C3. Fluorophores were diluted at 1:750 in TSA buffer: C1 was in 488; Channel 2 was in Cy3; Channel 3; Cy5. Once the procedure was completed, slides were counterstained with room temperature Prolong Gold Antifade Mountant with DAPI (Thermo Fisher Scientific, P36935).

Slides were scanned in a Zeiss Axioscan 7 microscope slide scanner with a 40x objective quadruple bandpass filter set with DAPI focus. CZI files were converted to 8-bit TIFF images in FIJI and appropriate brain regions (cortex, thalamus, hippocampus, and striatum) were outlined using the freehand selection tool. To assess levels of *Akap11*, *Prkaca*, and *Prka1a*, the mean intensity was quantified for each outlined brain region with background subtraction using the declive (VI), molecular layer. Four sections per animal were averaged and normalized to the WT to minimize batch effects.

### Viruses

FLIM-AKAR and FLIM-AKAR*^T^*^391*A*^ viruses were packaged in-house into a AAV2/AAV9 vector using plasmids that were available from Addgene (no. 60445 and 60446). The final viral constructs and concentration used in this study were: AAV2/1-FLEX-FLIM-AKAR (8.53 × 10^12^gc/ml), AAV9-FLEX-FLIM-AKAR (2.14 × 10^13^ gc/ml), AAV2-FLEX-FLIM-AKAR*^T^*^391*A*^ (1.23 x 10^13 gc/ml), AAV9-FLEX-FLIM-AKAR*^T^*^391*A*^ (7.67 × 10^12^ gc/ml). AAV2rg-Ef1a-Cre (1.39 × 10^13^ gc/ml) was obtained from Salk institute. While both AAV2 and AAV9 versions of the FLIM-AKAR and FLIM-AKAR*^T^*^391*A*^ were used in this study, we saw no noticeable difference in their expression level and baseline fluorescence lifetime when expressed in wild-type mice. During the experiments, FLIM-AKAR or mutant FLIM-AKAR*^T^*^391*A*^ virus was mixed with the Cre virus in a 4:1 ratio. A final volume of 250 nL was used for intracranial injection.

### Stereotaxic Surgery and Virus Injection

Mice were anesthetized with isoflurane and underwent bilateral injection of the virus mix (FLIM-AKAR + Cre or FLIM-AKAR*^T^*^391*A*^ + Cre). Typically, one side of the brain is injected with FLIM-AKAR + Cre, while the other side is injected with FLIM-AKAR*^T^*^391*A*^ + Cre while counterbalancing the sides of the brain, with the exception of twelve mice (mixed genotypes) injected with Dlight 3.8 virus instead of FLIM-AKAR*^T^*^391*A*^ + Cre (data not included). Virus injection was conducted using coordinates for the nucleus accumbens lateral shell (AP +1.20, ML ±1.50 from bregma, DV -4.2 from brain surface). Optical fibers (MFC_200/230-0.37_4.5mm_MF1.25_FLT from Doris Lenses) were placed bilaterally and ∼ 0.3 mm above the injection site.

### Fluorescence lifetime photometry (FLiP)

Thirty-three 12-20 week-old *Akap11* mutant mice of mixed genotypes were used for stereotaxic surgeries and subsequent FLiP measurements, including baseline, D1R antagonist and D2R antagonist expriments. Approximately equal number of male and female mice were used (n=10 WT, n=11 *Akap11^+/-^*and n=10 *Akap11^-/-^* ). The fluorescence lifetime and intensity of the FILM-AKAR (or FLIM-AKAR*^T^*^391*A*^) was measured in real time at 10kHz using a Fluorescent lifetime photometry system^106^. Customized MATLAB software was used to collect the lifetime measurement, and the lifetime data was down sampled to 10 Hz using a rolling mean. This setup has been described in detail with analysis code provided within^137^.

Animals were habituated into the recording room for at least 30 mins before the experiments. Baseline FLIM-AKAR lifetime was measured for 20 mins during the first measurement, and were then collected for 10 mins each during subsequent measurements, while animals were allowed to freely move in an open field (18x18 cm). Drug injections (dopamine D1/D2 antagonists) during pharmacology experiments were conducted after 10 mins baseline lifetime measurement, and the lifetime data were collected for at least another 20 mins following intraperitoneal injections.

The drugs used in our study include: D1R antagonist (SKF 83566 hydrobromide, Cayman Chemical #29480; 3 mg kg^−1^), D2R antagonist (eticlopride hydrochloride, Cayman Chemical #29112; 0.5 mg kg^−1^). Drugs were dissolved in sterile saline at appropriate concentrations and were injected at 0.1 ml solution per 10 g of mouse. An average injection including scruffing the mice took ∼30 s and animals were habituated to the injection procedure with sterile saline twice before the pharmacology experiments.

The experimenter was blinded to the genotype of the mice during the experiments and subsequent data analysis. Two out of thirty-three FLIM-AKAR measurements were excluded from the study due to the low expression of the FILM-AKAR causing the autofluorescent contribution to the recorded lifetime to be greater than 20% based on our estimation. The exclusion criteria was determined before the experiment while the experimenter was still blinded to the mice genotypes.

### Post- FLiP immunofluorescence expression validation

After the FLiP experiments were completed, all mice were perfused transcardially with PFA. Coronal brain sections (80μm) near the optical fiber implants were sliced using a vibratome (Leica VT1200S). The brain slices were mounted with Prolong Gold Antifade Mountant with DAPI (Thermo Fisher Scientific, P36935) and native fluorescence of FLIM-AKAR was imaged with a Zeiss Axioscan 7 slidescanner.

To assess subcellular localization of FLIM-AKAR, additional brain slices were imaged with a Zeiss 900 LSM confocal microscope. Images were acquired at magnifications of 20x with the green-channel laser power set to 0.2 mW. Z-stack images were taken at 63x under oil with a step size of 0.3 um and averaging was set to 4x. Image regions were matched across animals based on coordinates determined on proximity to bregma (y = 1.54 mm) and site of the injection. To facilitate clear visualization of the FLIM-AKAR sensor’s subcellular localization, images were specifically taken in areas of sparse viral expression located towards the lateral edge of the nucleus accumbens core. Schematic diagrams in Supplementary Figures 11b,c are adapted with permission^138^.

### Amphetamine induced-hyperlocomotion (AIH)

*Akap11* mice that were 10-12 weeks old with roughly equal numbers of male and female mice were used for the AIH experiment. n=18 WT, n=22 *Akap11^+/-^* and n=14 *Akap11^-/-^* were used for this experiment. Mice in each genotype were randomly assigned to the saline or amphetamine groups. Before the experiment, dextroamphetamine sulfate (Millipore Sigma 1180004; salt corrected) was dissolved in water at 0.2667 mg/mL and was pH corrected to 7.3. Mice were weighted and were allowed to habituate in the behavioral room (under white light) for 30 mins before the start of the experiment. For the measurement of locomotion, mice were placed in Omnitech Electronics Inc SuperFlex Open Field apparatus (Omnitech Electronics Inc), which was cleaned with 70% ethanol between each experiment. The Fusion software (Omnitech Electronics Inc) was used for data collection and analysis. First, a 90 min baseline locomotion was recorded, followed by subcutaneous injection of amphetamine at a dose of 2mg/kg or saline.

Following drug injection, the locomotion data were collected for another 90 min. Locomotion data were measured by infrared beams and were measured by beam breaks, and were grouped into 5-min bins for analysis.

### Statistics

Statistics for immunoblots, fluorescence lifetime photometry (FLiP), and AIH were calculated in GraphPad Prism, with each dot representing an individual animal. Immunoblots and FLiP experiments including the genotype factor and a second factor (Age, Brain region, Protein interactor, or FLIM-AKAR variant) were quantified with two-way ANOVA. For all analyses with a significant main effect of genotype, post hoc Tukey’s multiple comparisons were conducted for all possible comparisons within the same level of the second factor (e.g. WT, *Akap11^+/-^*and *Akap11^-/-^* were compared within one age, but not between different ages). For Supplementary Figs. 3b,c and 4g, where *Akap11^-/-^*was excluded, Sidak’s multiple comparisons test was used. For immunoblot, and FLiP experiments where genotype was the only variable, statistically analysis were conducted by one-way ANOVA followed by Tukey’s post hoc test to compare all genotype pairs. Further statistical information is available in Supplementary Data 1.

## Data availability

The original mass spectra and the protein sequence database used for searches have been deposited in the public proteomics repository MassIVE (http://massive.ucsd.edu) and are accessible at ftp://MSV000095852@massive.ucsd.edu. These datasets will be made public upon acceptance of the manuscript. Processed and raw bulk and single nucleus RNA sequencing data have been deposited at Gene Expression Omnibus (GEO) with the following IDs: bulk RNAseq: GSE306677. single nucleus RNAseq: GSE272098. For analyses of *Akap11* transcript abundance in wild-type mice, the following publicly available datasets were analyzed: GSE218573. All software used in the analysis are publicly available. Source data are provided with this paper.

## Code availability

No specialized code was developed for this manuscript. Software and packages that were utilized for analysis are publicly available and referenced throughout the methods.

## Acknowledgements

We thank Kris Dickson and members of the Sheng and Granger Labs for valuable input; Ben Neale, Tarjinder Singh and Calwing Liao for consultations on human genetic evidence; Kira Brenner and Jacalyn Fahey for assistance with immunohistochemistry; Abraham Eafa and Miriam Kpoyto for help with colony management; Miriam Kpoyto for help with amphetamine-induced hyperlocomotion experiments; Abe Eafa and Nate Hodgson for help with behavioral assays; Guido Gottardo, Reivi Voci, Mara Rigamonti and Tecniplast for assistance with the homecage recording system; Michael Dolan and Chuck Vanderburg for guidance on transcriptome experiments; Michael Sherman for packaging AAVs; Tyler Caron and Matthew Beck for help with mouse colony management and *in vivo* experiment logistics; the Broad Institute Genomic Platform for assistance with RNA sequencing. Artwork of a saggital mouse brain was adapted from open access artwork created and shared by Federico Claudi (doi:10.5281/ZENODO.3925911). We thank the IDDRC Animal Behavior and Physiology Core, funded by NIH/NICHD P50 HD105351

## Author contributions

B.J.S. and M.S. conceived and designed the -omics, biochemical, and behavioral experiments. Y.G., M.S. and A.G. conceived and designed fiber photometry, electrophysiology, pharmacological and behavioral experiments. B.J.S. and M.S. wrote the manuscript, with help from Y.G. and A.G. and input from all authors. B.J.S. and Y.G. performed the majority of experiments and analyses in this paper. Specifically, B.J.S. performed immunoprecipitations, immunoblots, immunohistochemistry, synapse purifications, nuclei isolation, RNAseq, homecage activity measurements and analyzed these data. Y.G. performed FLiP measurements, slice electrophysiology, dopamine receptor pharmacology, amphetamine-induced hyperlocomotion and analyzed these data; Y.G. analyzed behavioral data from the IDDRC animal behavior and physiology core. A.N. and K.P.M. processed the transcriptome data, including quality control, reference genome alignment, cell type clustering, annotation and differential expression analysis, with guidance from S.K.S. and supervision from J.Z.L. M.J.K. performed nuclei isolation and library preparation for 12 week striatum and PFC snSeq datasets. B.L. designed the FLiP recording set up, calibrated the measurements and developed the analysis pipeline with guidance from B.L.S. K.B. performed LC-MS/MS for synapse proteomics and synapse phosphoproteomics and A.V.T. performed LC-MS/MS for immunoprecipitations with guidance from H.K. and S.A.C and analysis help from D.R.M. J.A. conducted motion sequencing tests and D.M. analyzed these data. N.H. conducted associative learning and sensory discrimination assays, with help from J.T. and supervision from Z. Fu. J.A. and C.G. helped with immunoblots. S.N. performed smFISH. I.P. and N.S. assisted with brain tissue collection and immunohistochemistry and N.S. additionally designed and performed rotarod behavioral tests. H.P. and N.M.N. assisted with nuclei isolation and snSeq library preparation with supervision from E.Z.M. E.Z.M. designed the snSeq protocol and guided snSeq analysis. S.A., Z. Farsi and X.L. helped with data analysis. A.H. conducted confocal microscopy with supervision from P.S.K. N.M. conducted phosphoproteomics motif analysis with supervision from B.S. B.D. purified synapse fractions for proteomics/phosphoproteomics and provided guidance and supervision for biochemical experiments. M.S. and A.G. supervised the project and acquired funding. This work was supported by the Stanley Center for Psychiatric Research and the Harvard Brain Science Initiative Bipolar Disorder Seed Grant.

## Competing Interests

M.S. is cofounder and scientific advisory board (SAB) member of Neumora Therapeutics and serves on the SAB of Biogen, Proximity Therapeutics and Illimis Therapeutics. S.A.C. is a member of the SAB of Kymera, PTM BioLabs, Seer, and PrognomIQ. B.S. serves on the Scientific Advisory Board of Annexon Biosciences and is a minor shareholder of Annexon. B.D. is a full-time employee of Vigil Neuroscience. E.Z.M. is an academic founder of Curio Bioscience. The other authors declare no competing interests.

---

**Supplementary Figure 1.**
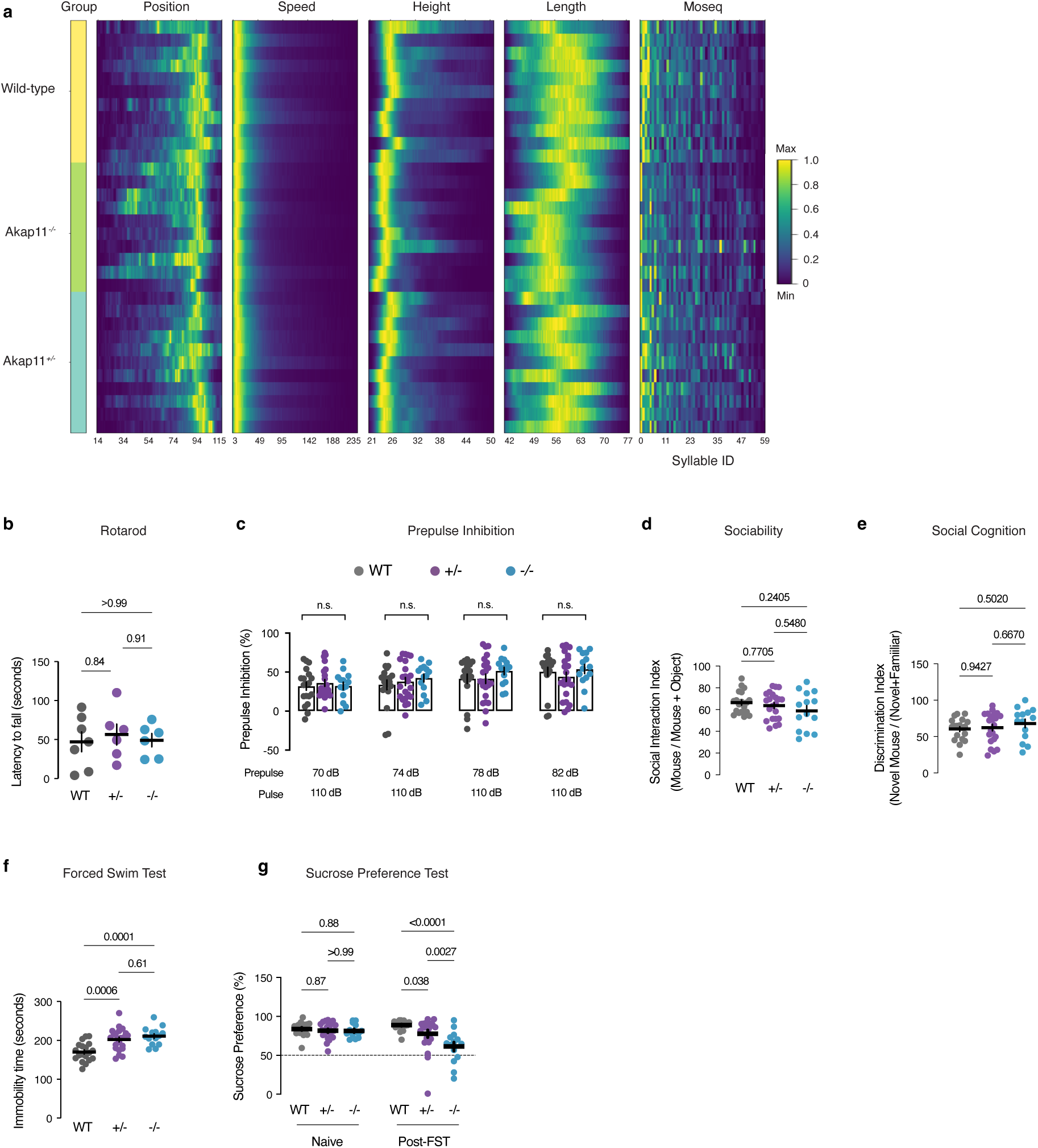
Behavioral survey of *Akap11* mutant mice, related to Fig. 1. **a**, Heatmaps showing the position, speed, height, length, or MoSeq (syllable usage) by each mouse, grouped into WT (top), *Akap11^+/-^* (bottom), and *Akap11^-/-^* (middle). **b**, Average latency to fall across 4 rotarod test trials. **c**, Prepulse inhibition to an acoustic stimulus, measured as a percentage of accelerometer response to the test pulse, relative to a less intense prepulse. **d**, Performances of WT and *Akap11* mutant mice during the three-chamber test for sociability. Social interaction index is defined as: time(mouse)/(time(mouse) + time(object)) x 100%. **e**, Performances of WT and *Akap11* mutant mice during the three-chamber test for social cognition. Discrimination index is defined as: time(novel)/time(novel) + time(familiar) x 100%. **f**, Immobility duration during the forced swim test (FST). **g**, Sucrose preference of *Akap11* mutant and WT animals before and after undergoing FST. **b-g**, data are represented as mean +/- SEM **b,d,e,f,** One-way ANOVA with Tukey’s post hoc test. See Supplementary Data 1 for statistics. **c,g,** Repeated measures two-way ANOVA with Tukey’s post hoc test. See Supplementary Data 1 for statistics.

**Supplementary Figure 2:**
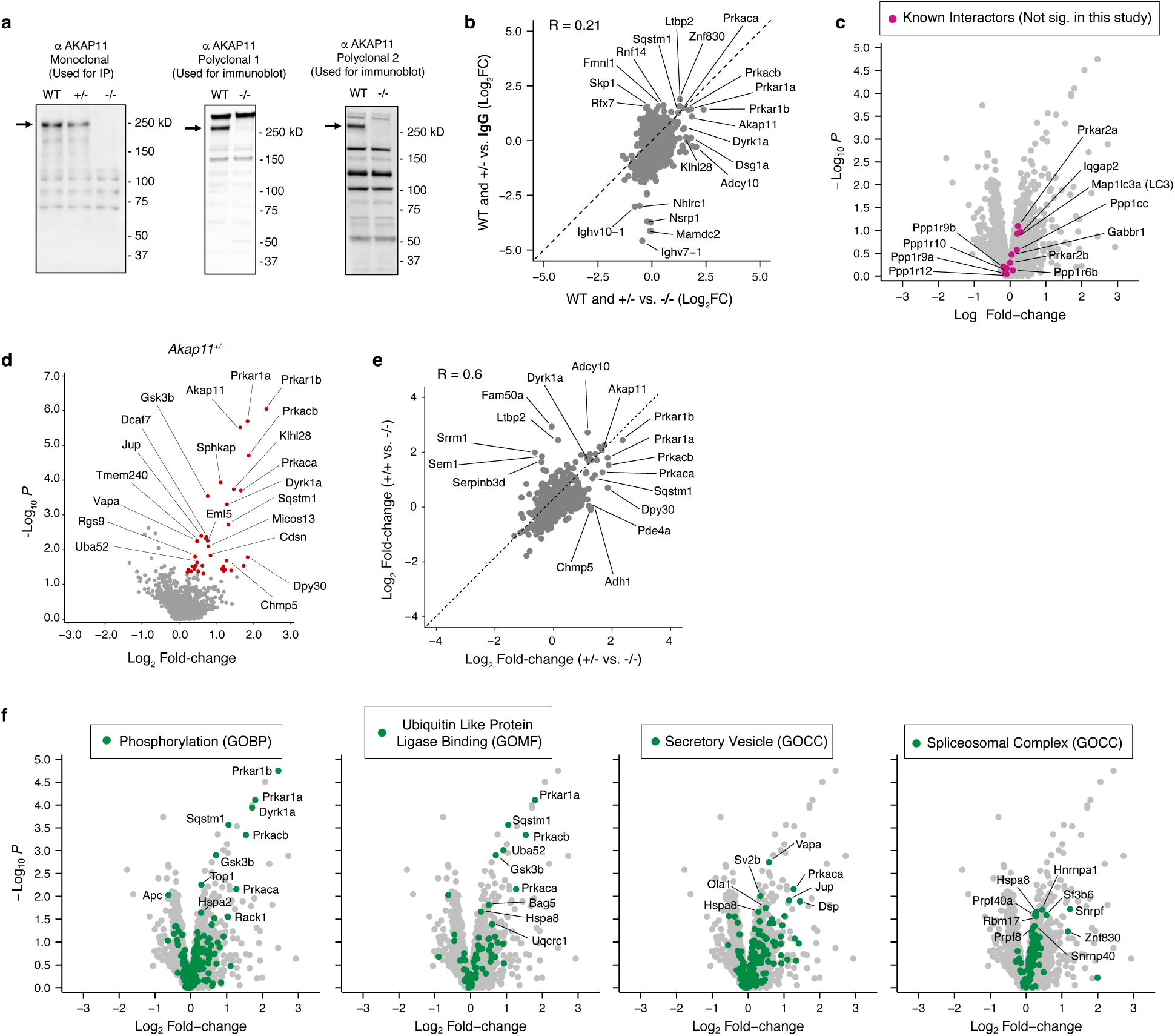
AKAP11-interacting proteins and pathways in WT and in *Akap11^+/-^*, related to Fig. 2. **a,** Immunoblots of mouse cortex total lysate using AKAP11 antibodies used in this study. **b,** Log2FC scatterplot comparing *Akap11^-/-^* vs. IgG as controls for AKAP11 IP. In these data, AKAP11 IP in WT and *Akap11^+/-^* animals were compared together against AKAP11 IP in *Akap11^-/-^* animals or IgG in WT animals. **c,** Volcano plot of AKAP11 CoIP-MS (WT vs. *Akap11^-/-^*). Highlighted in magenta are proteins that have been previously reported to interact with AKAP11, but were not significantly enriched in this study. **d,** Volcano plot of AKAP11 coIP-MS, comparing IP in *Akap11^+/-^* vs. *Akap11^-/-^* cortex. Proteins with nominal *P* value < 0.05 and Log2FC > 0 are colored red. **e,** Log2FC scatterplots comparing all proteins detected by AKAP11 IP in WT versus *Akap11^+/-^*. **f,** Volcano plot of AKAP11 coIP-MS. Highlighted in green are proteins within gene ontology (GO) pathways that were significantly found to be enriched by GSEA. (GOBP, Gene Ontology Biological Process; GOCC, Gene Ontology Cellular Component; GOMF, Gene Ontology Molecular Function). **b,e,** R = Pearson correlation.

**Supplementary Figure 3.**
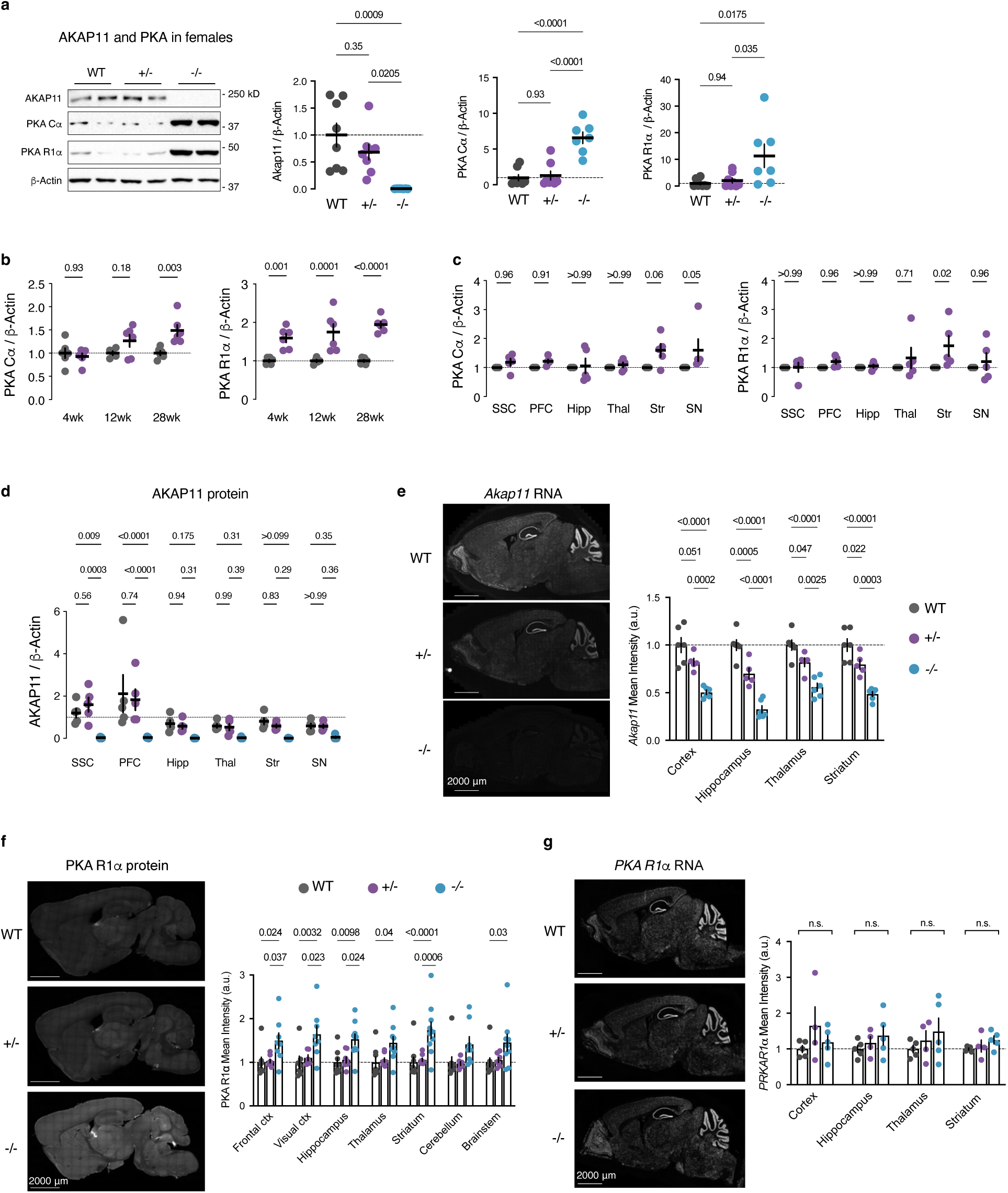
PKA protein levels in *Akap11^+/-^* and *in situ*, related to Fig. 3. **a,** Quantifications of AKAP11 and PKA subunit protein levels in female 12wk WT and *Akap11^+/-^* total lysate. One-way ANOVA with Tukey’s post hoc test. **b,c,** Quantifications of PKA subunit proteins levels in WT and *Akap11^+/-^* total lysate. These data are also shown in Fig. 3b,c, but are rescaled here without *Akap11^-/-^*for clarity. Two-way ANOVA with Sidak’s post hoc test. **d,** Quantification of immunoblots from microdissected tissue total lysate from 4wk mice using antibodies against AKAP11. Representative immunoblots for these measurements are shown in Fig. 3c. *P*-values are calculated without *Akap11^-/-^* in multiple comparisons. Unlike other figures, these data are not normalized to WT within a brain region, in order to display differences in AKAP11 protein across brain regions. **e,** *In situ* hybridization measurements of *Akap11* RNA. Right, quantification of mean intensity measurements within anatomical subregions. Measurements are normalized to WT controls within a region. **f,** Immunofluorescence measurements of PKA R1α protein *in situ*. Right, quantification of mean PKA R1α immunofluorescence intensity measurements within anatomical subregions. Multiple comparisons adjusted *P* values < 0.05 are displayed above. **g,** *In situ* hybridization measurements of *PKA R1α* RNA. Right, quantification of mean intensity measurements within anatomical subregions. Measurements are normalized to WT controls within a region. **d-g,** Two-way ANOVA with Tukey’s post hoc test. See Supplementary Data 1 for statistics. **a-g,** Data are represented as mean +/- SEM.

**Supplementary Figure 4.**
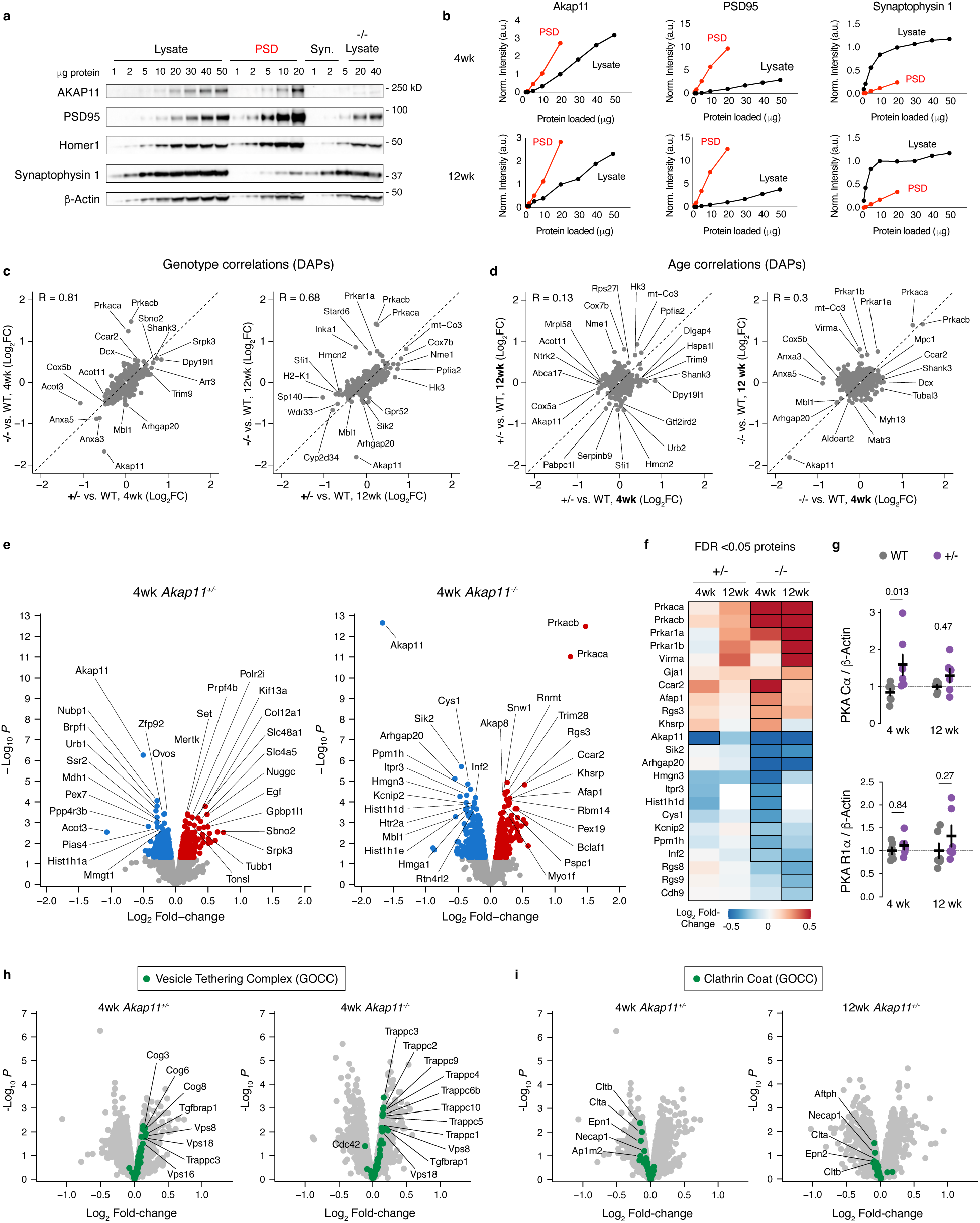
Comparisons of juvenile and adult synapse proteomes, related to Fig. 4. **a**, Immunoblots of AKAP11 and synapse proteins across different amounts of lysate, synaptosome, and PSD-enriched fractions of 12wk cortex. **b,** Quantifications of immunoblots in (a). Values are normalized to 20μg lysate. **c,** Log_2_FC scatter plots comparing DAPs in *Akap11^+/-^* vs. *Akap11^-/-^* synapse proteomes. **d,** Log_2_FC scatter plots comparing DAPs in 4wk vs. 12wk synapse proteomes. **e,** Volcano plots of 4wk synapse proteomics data. Upregulated and downregulated DAPs are colored red and blue, respectively. **f,** Heatmap of adjusted *P*-value (FDR) < 0.05 proteins across 4wk and 12wk synapse proteomes. Black border indicates FDR significance, including corrections for comparisons between *Akap11^+/-^* and *Akap11^-/-^*. **g,** Quantifications of PKA subunit protein levels in WT and *Akap11^+/-^* synapse fractions. These data are also shown in Fig. 4c, but are rescaled here without *Akap11^-/-^* for clarity. Data are represented as mean +/- SEM. Two-way ANOVA with Sidak’s post hoc test. See Supplementary Data 1 for statistics. **h,** Volcano plots of 4wk synapse proteomics data in *Akap11^+/-^*. Proteins within the Vesicle Tethering Complex gene set are highlighted in green. **i,** Volcano plots of synapse proteomics data in *Akap11^+/-^*. Proteins within the “Clathrin Coat” gene set are highlighted in green for 4wk and 12wk animals.

**Supplementary Figure 5:**
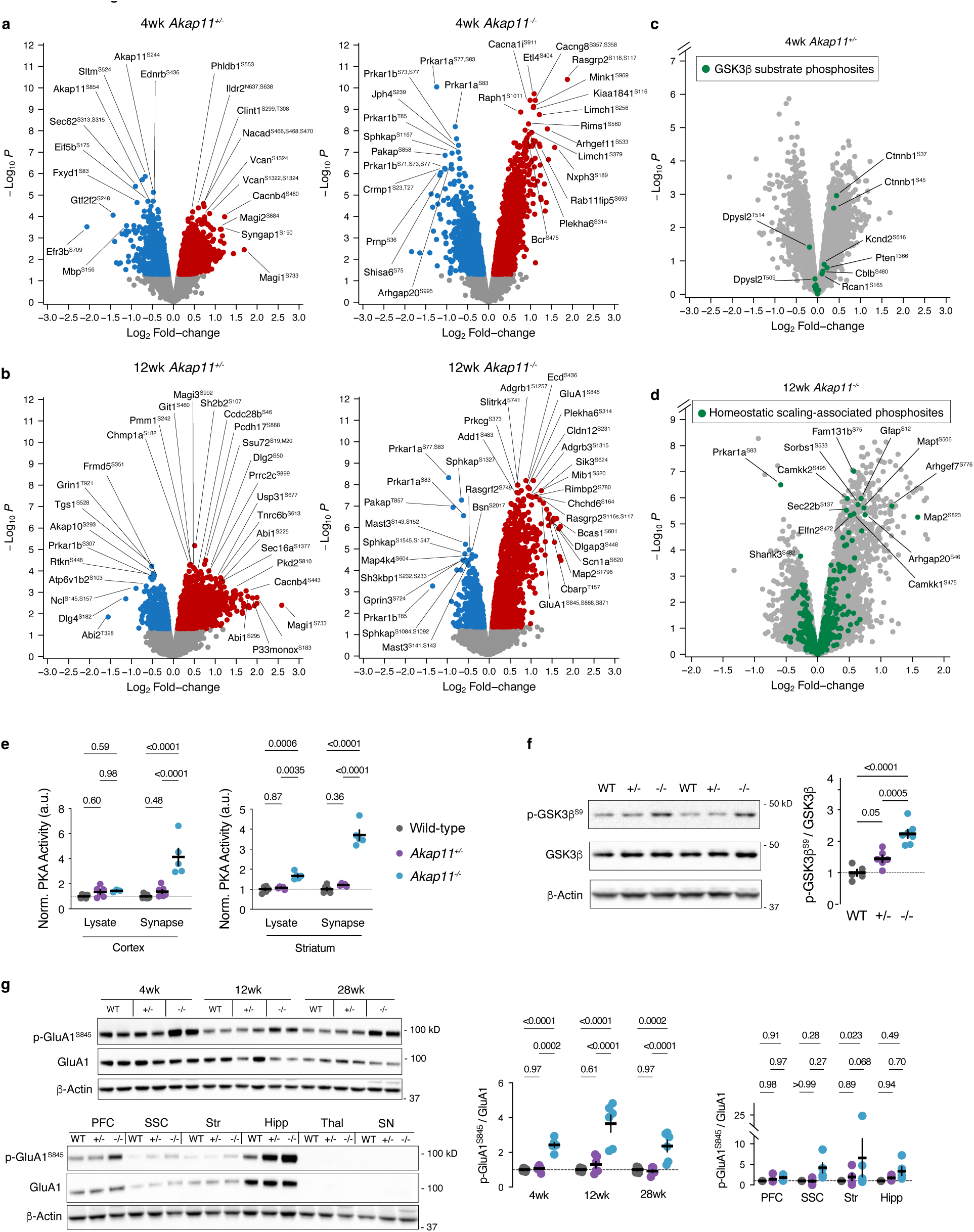
Phosphoproteomic correlates of synaptic plasticity, related to Fig. 5. **a,** Volcano plots of synapse phosphoproteomics in 4wk *Akap11* mutants relative to WT. Upregulated and downregulated DAPPs are colored red and blue respectively. AKAP11 phosphopeptides are omitted from *Akap11^-/-^* plots, for clarity. **b,** Volcano plots of synapse phosphoproteomics in 12wk *Akap11* mutants relative to WT. Upregulated and downregulated DAPPs are colored red and blue respectively. AKAP11 phosphopeptides are omitted from *Akap11^-/-^* plots, for clarity. **c,** Volcano plots of 4wk synapse phosphoproteomics for *Akap11^+/-^*. Known GSK3β substrates from PhosphoSitePlus are labeled in green. **d,** Volcano plots of 12wk synapse phosphoproteomics for *Akap11^+/-^*. Phosphosites that are altered during scaling (24 hours of bicuculline or tetrodotoxin treatment, FDR <0.05 in Desch et al., 2021)^72^ are labeled in green. **e,** ELISA-based PKA activity measurements comparing total lysate and synapse fractions of cortex (left) and striatum-enriched tissue (right). These measurements appear in Fig. 5e, but have been normalized here to wild-type for more clear representation of fold-change. Two-way ANOVA with Tukey’s post hoc test. **f,** Immunoblotting and quantification of p-GSK3β^S9^ and GSK3β in 4wk cortical total lysate. One-way ANOVA with Tukey’s post hoc test. See Supplementary Data 1 for statistics. **g,** Immunoblotting and quantification of p-GluA1^S845^ and GluA1 in 4wk, 12wk or 28wk cortical or in microdissected brain region total lysate. Two-way ANOVA with Tukey’s post hoc test. See Supplementary Data 1 for statistics.

**Supplementary Figure 6:**
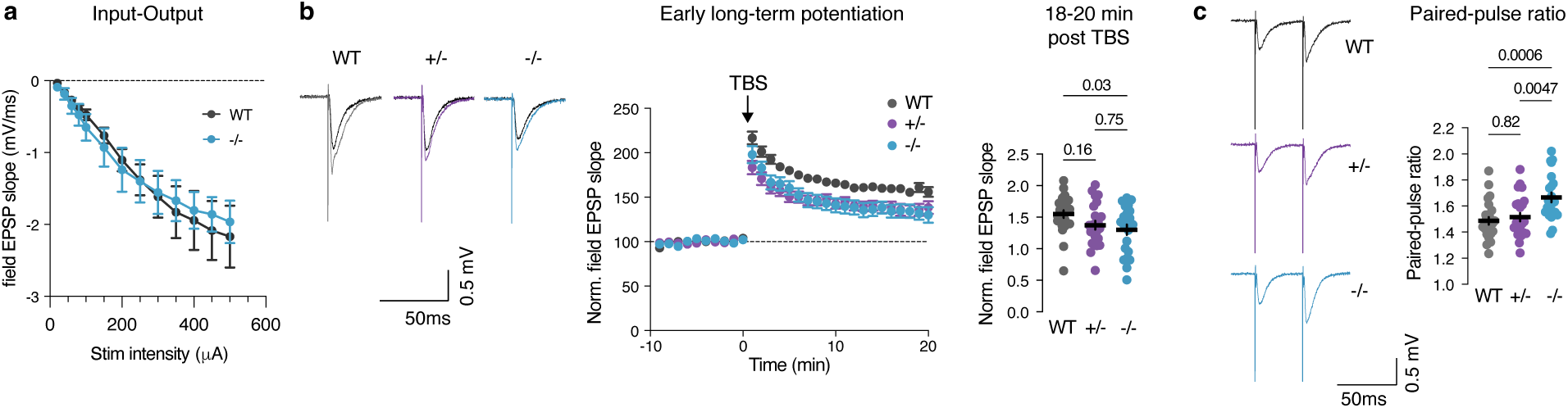
Electrophysiological characterization of synaptic strength and plasticity, related to Fig. 5. **a,** Input-output (I-V) curves of field excitatory postsynaptic potential (EPSP) responses evoked with varying stimulation intensities (20-500µA). **b,** Left, representative traces of field EPSPs before and after LTP induced by theta burst stimulation (TBS). Traces were generated by averaging the data points during the first (baseline; light lines) and last (post LTP; dark lines) two mins of the experiment. Middle, quantification of field EPSP slopes over the 30-min long early LTP experiment. Each dot represents the average field EPSP slope over 1 min. Right, quantification of field EPSP slope in minutes 18-20 following TBS. **c,** Left, representative traces of paired pulse responses at 40ms stimulation intervals. Right, quantification of paired pulse ratios from *Akap11* mutants and WT brain slices. **a-c,** Data are represented as mean +/- SEM. One-way ANOVA with Tukey’s post hoc test. See Supplementary Data 1 for statistics.

**Supplementary Figure 7:**
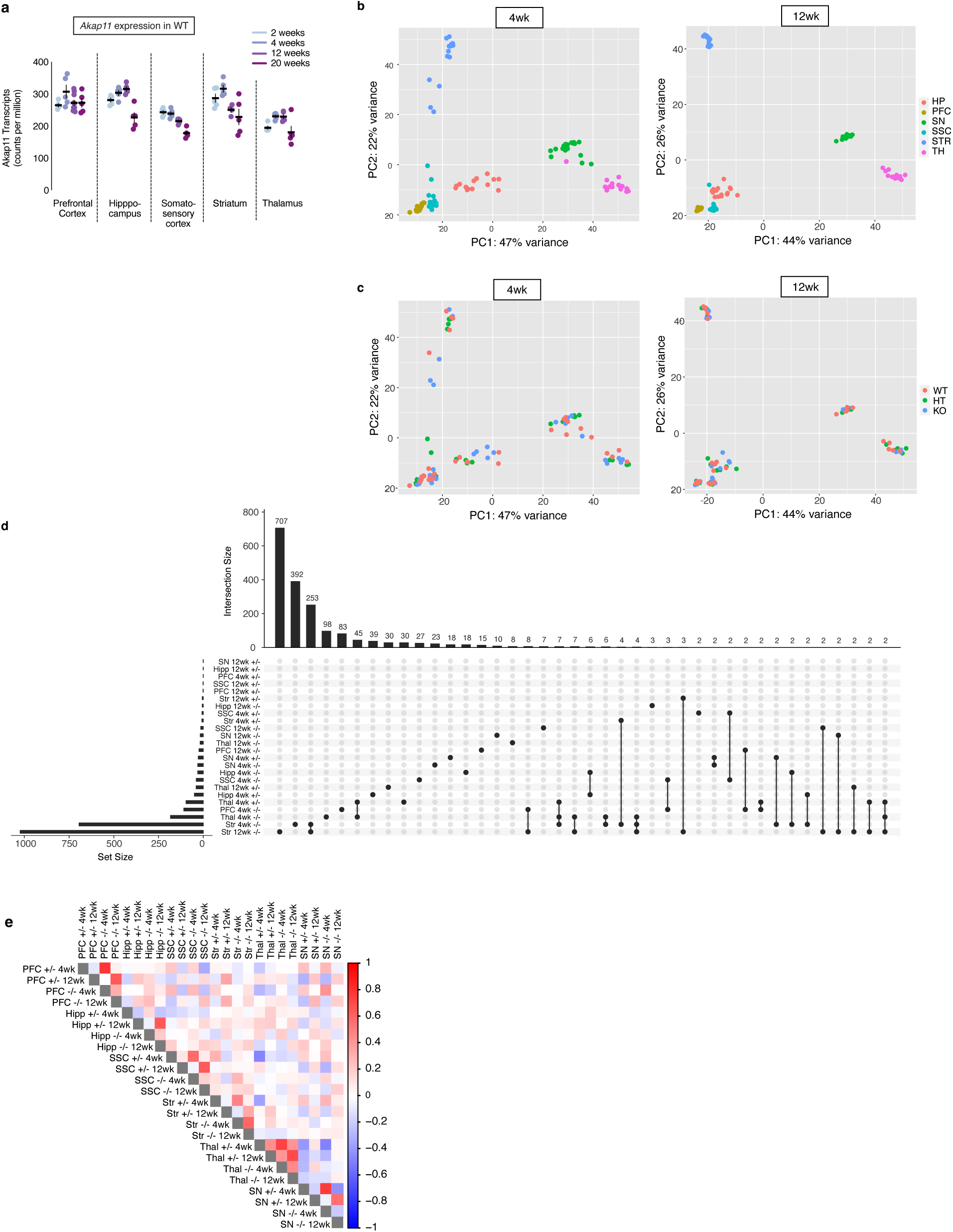
Comparisons between brain regions, ages, and genotypes in bulk RNA sequencing, related to Fig. 6. **a,** *Akap11* mRNA expression across brain regions and ages in wild-type mice, analyzed from data published in Farsi et al., 2023^12^. Data are represented as mean +/- SEM. **b,** PCA plots of bulk RNA sequencing experiments, colored by brain region. **c,** PCA plots of bulk RNA sequencing experiments, colored by genotype. **d,** Upset plot showing overlapping and unique DEGs (FDR < 0.05) across brain regions, ages and genotypes in bulk RNA sequencing experiments. **e,** Log_2_FC correlations between nominally significant genes (*P* < 0.05) across brain regions, ages, and genotypes. Color indicates Spearman correlation values.

**Supplementary Figure 8:**
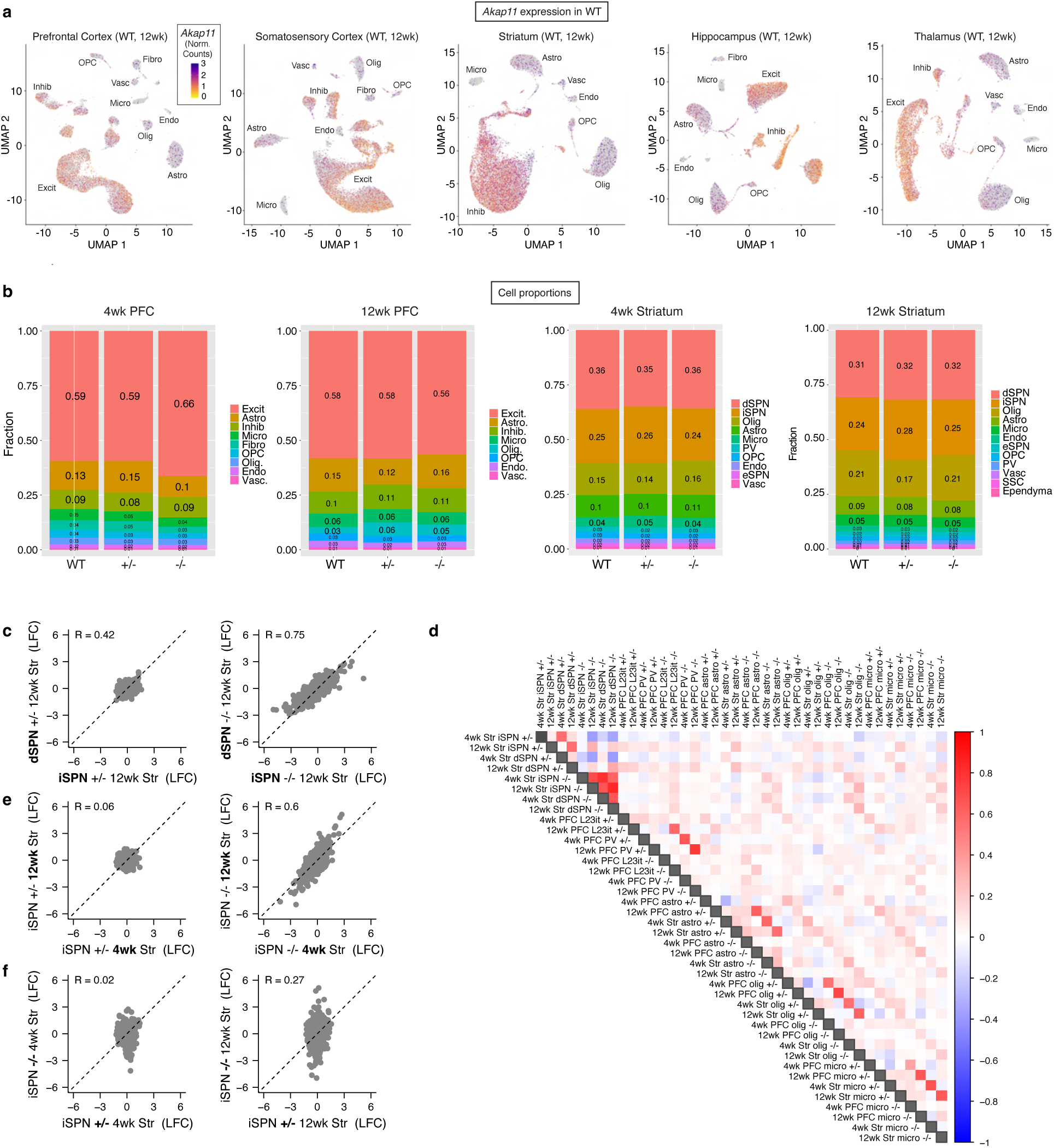
Comparisons between brain regions, age and cell types in snRNA-seq, related to Fig. 6. **a,** *Akap11* expression across major cell types on uniform manifold approximation and projection (UMAP) representations of cell types in different brain regions and ages of wild-type mice. These data appears in Farsi et al., 2023^12^. **b,** Fraction of each cell type identified across genotypes. **c,** Log_2_FC scatterplots comparing all genes for iSPN vs dSPN subtypes. **d,** Log_2_FC correlations between nominally significant genes (*P* < 0.05) across cell types. Color indicates Pearson correlation values. **e,** Log_2_FC scatterplots comparing all genes at 4wk vs. 12wk, in iSPNs. **f,** Log_2_FC scatterplots comparing all genes in *Akap11^+/-^* vs. *Akap11^-/-^*, in iSPNs.

**Supplementary Figure 9:**
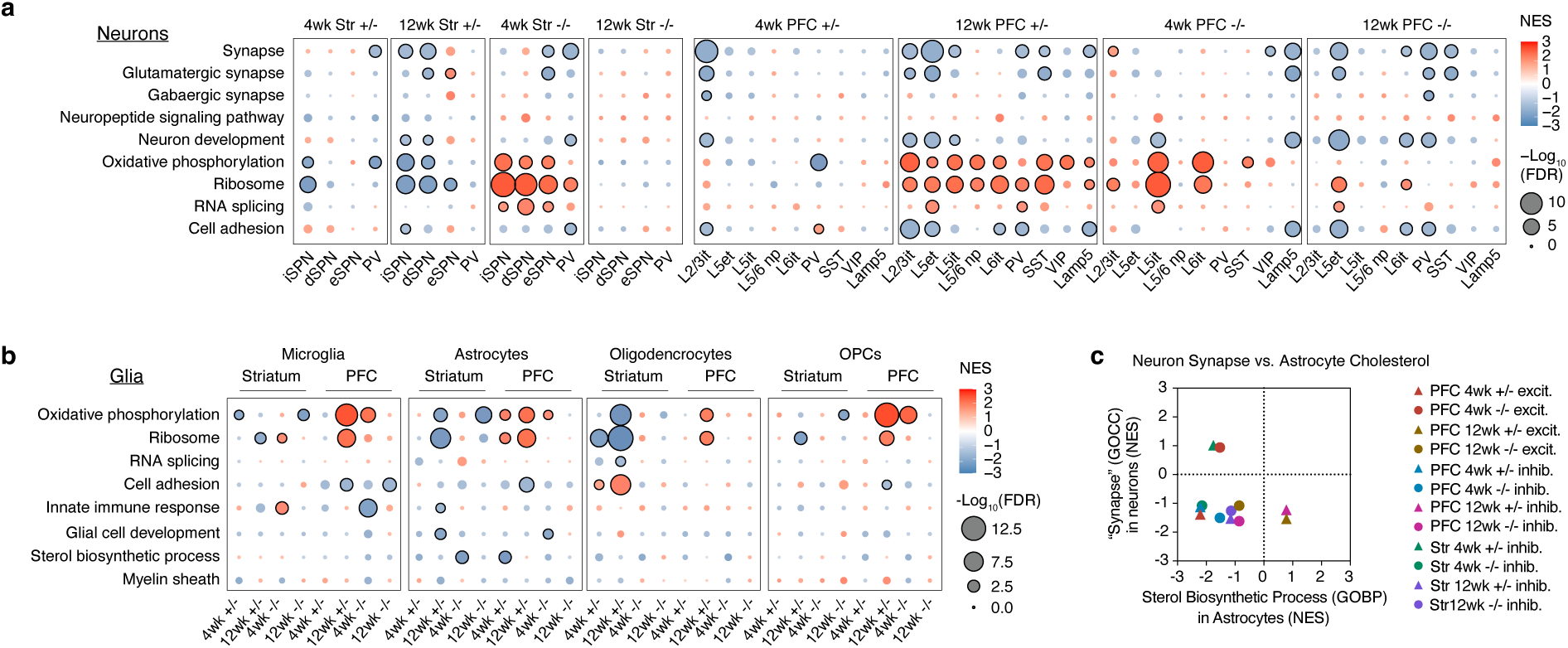
Neuronal-synapse and astrocyte-cholesterol related transcriptomic changes detected by snRNA-seq, related to Fig. 6. **a,** GSEA from neurons identified by snRNA-seq of striatum and PFC, showing MsigDB pathways of interest. **b,** GSEA results from glia snRNA-Seq, showing, showing MsigDB pathways of interest, including pathways related to glia function. **c,** Comparison of NES from synapse pathways in neurons vs. cholesterol biosynthesis pathways in astrocytes.

**Supplementary Figure 10:**
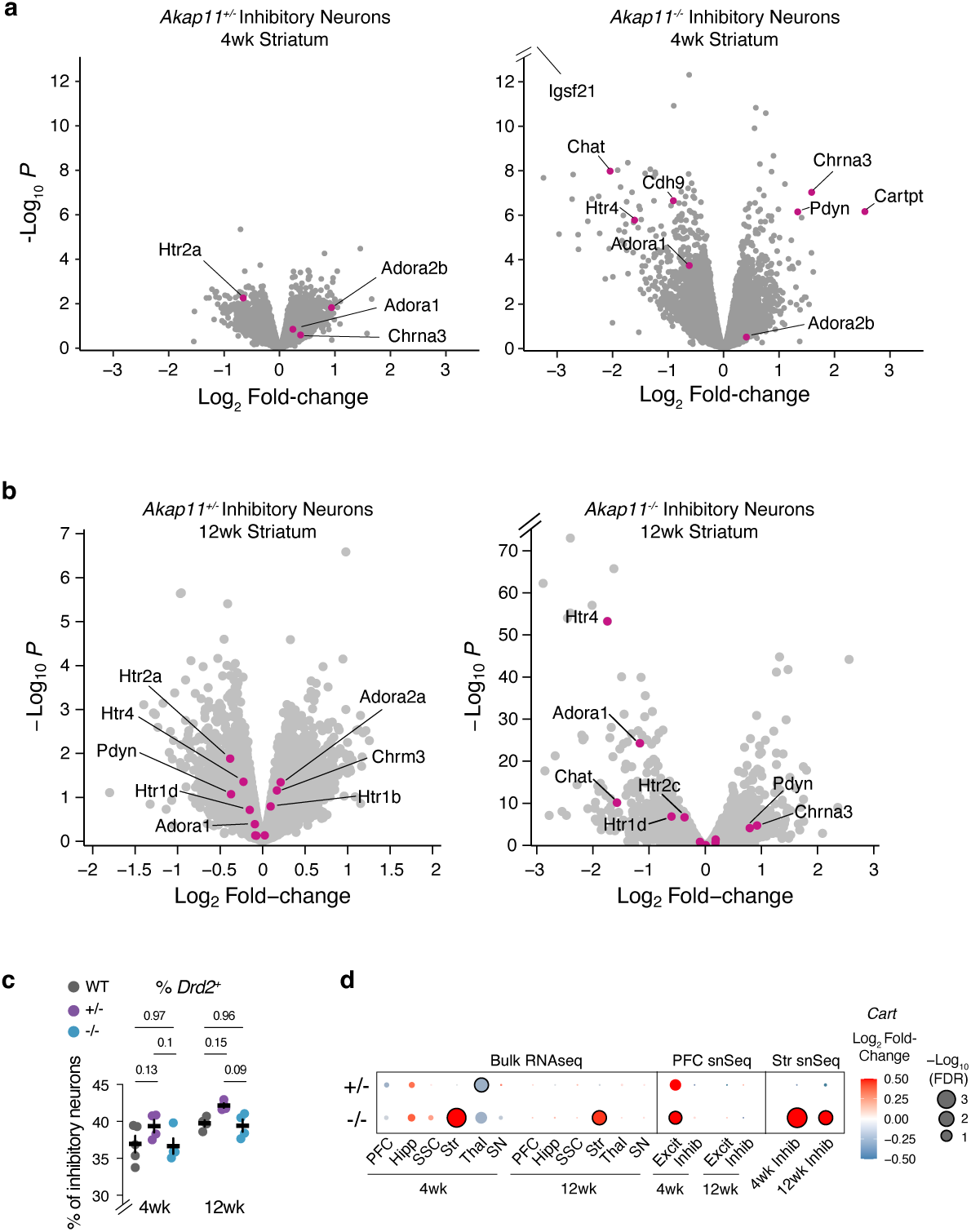
Evidence for transcriptomic alteration of neuromodulation-related genes, related to Fig. 6. **a,** Volcano plots of transcriptomic changes identified by snRNA-seq analysis in 4wk striatal inhibitory neurons. Genes of interest related to neuromodulation and circuit development are highlighted in magenta. **b,** Volcano plots of transcriptomic changes identified by snRNA-seq analysis in 12wk striatal inhibitory neurons. Genes of interest related to neuromodulation and circuit development are highlighted in magenta. **c,** Percentage of Drd2^+^ inhibitory neurons in the striatum. Data are represented as mean +/- SEM. Two-way ANOVA with Tukey’s post hoc test. **d,** Log_2_FC and FDR-adjusted *P*-values for *Cart* across bulk and snRNA-seq.

**Supplementary Figure 11:**
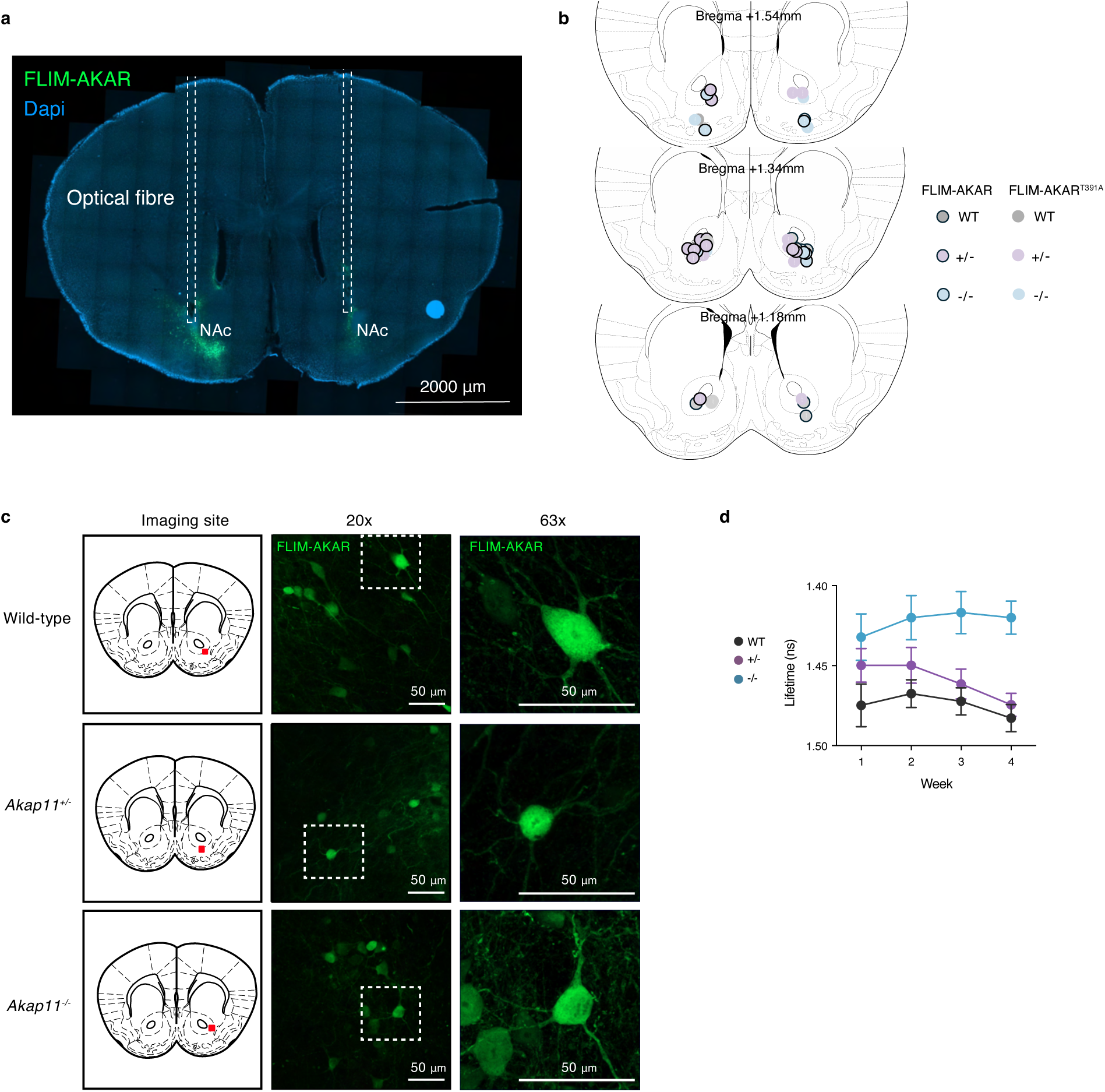
FLIM-AKAR injections sites and longitudinal measurement of PKA activity, related to Fig. 7. **a,** Immunohistochemical validation of AAV injection site targeting of NAc. **b,** Injection site targets of each animal in this study. Animals of different genotypes are differentiated by color, and FLIM-AKAR variants that were injected into contralateral hemispheres are differentiated by black border. **c,** Native fluorescence of FLIM-AKAR, imaged under a confocal microscope. Left, schematic of approximate imaging site. **d,** Fluorescence lifetime of FLIM-AKAR across 4 fiber photometry recording sessions across weeks.

**Supplementary Figure 12:**
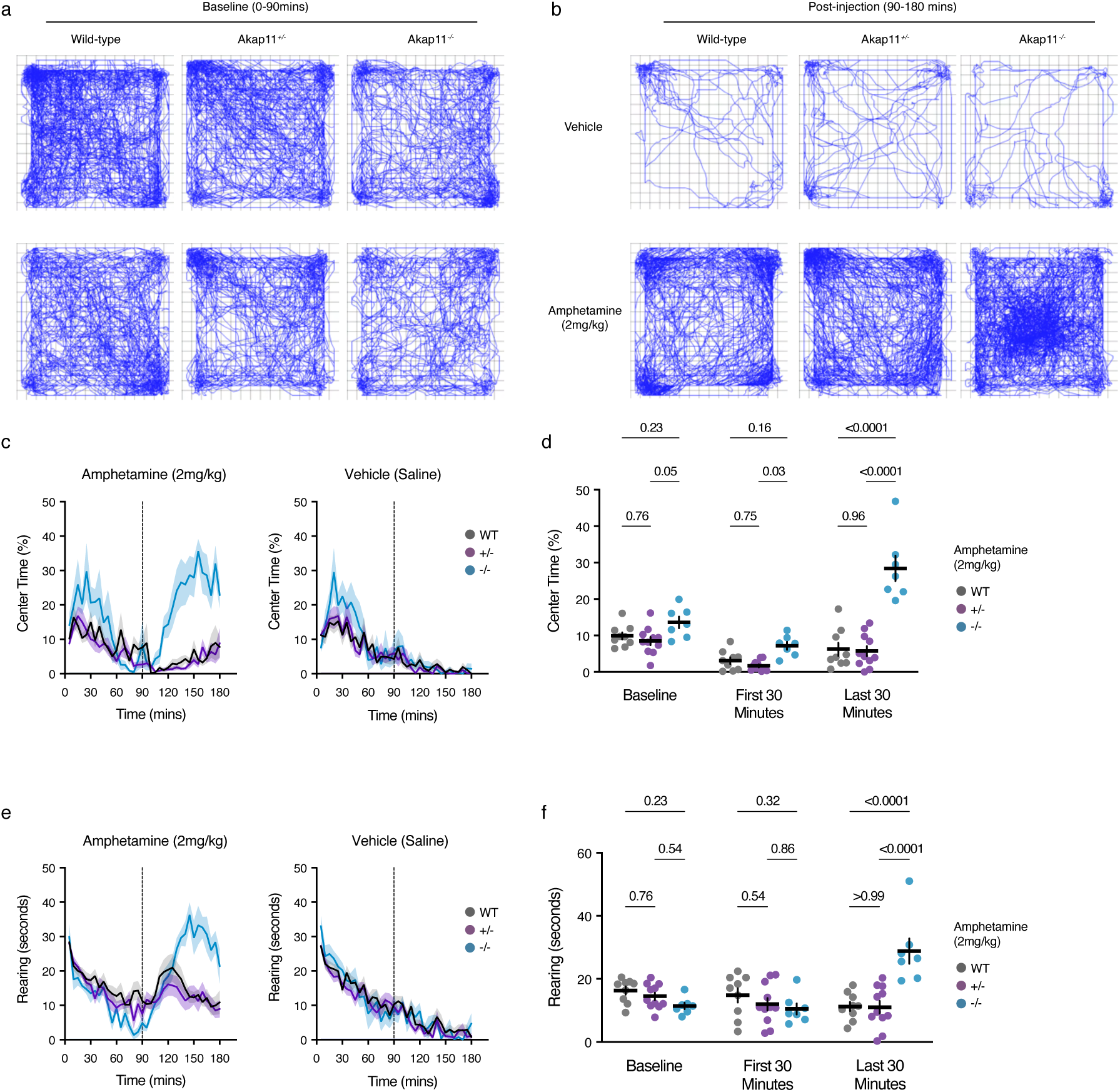
Amphetamine induces excessive open-field center time and rearing behaviors in *Akap11^-/-^* mice, related to Fig. 7. **a,b**, Representative activity traces of mice in the open field arena, before **(a)** and after **(b)** amphetamine (2 mg/kg) or saline administration. The same animals are shown in **(a)** and **(b)**, before and after vehicle (top rows) or amphetamine (bottom rows) administration. **c**, Centre time (%) after administration of amphetamine or saline to WT or *Akap11* mutants. **d**, Quantification of center time (%) during baseline (0-90mins), first 30 minutes post injection (90-120mins), and during the last 30 minutes (150-180mins) of the trial. Repeated measures two-way ANOVA with Tukey’s post hoc test. See Supplementary Data 1 for detailed statistics. **e**, Rearing time (s) after amphetamine (2 mg/kg) or saline administration to WT or *Akap11* mutants. **f**, Quantification of rearing time (s) during baseline (0-90mins), first 30 minutes post injection (90-120mins), and during the last 30 minutes (150-180mins) of the trial. Repeated measures two-way ANOVA with Tukey’s post hoc test. See Supplementary Data 1 for detailed statistics.

## Notes

### Summary of Updates

Added new data (Figure 1 - behavioral tests)

